# Synteny-guided resolution of gene trees clarifies the functional impact of whole genome duplications

**DOI:** 10.1101/2020.01.30.926915

**Authors:** Elise Parey, Alexandra Louis, Cédric Cabau, Yann Guiguen, Hugues Roest Crollius, Camille Berthelot

## Abstract

Whole genome duplications (WGD) have major impacts on the evolution of species, as they produce new gene copies contributing substantially to adaptation, isolation, phenotypic robustness, and evolvability. They result in large, complex gene families with recurrent gene losses in descendant species that sequence-based phylogenetic methods fail to reconstruct accurately. As a result, orthologs and paralogs are difficult to identify reliably in WGD-descended species, which hinders the exploration of functional consequences of WGDs. Here we present SCORPiOs, a novel method to reconstruct gene phylogenies in the context of a known WGD event. WGDs generate large duplicated syntenic regions, which SCORPiOs systematically leverages as a complement to sequence evolution to infer the evolutionary history of genes. We applied SCORPiOs to the 320-million-year-old WGD at the origin of teleost fish. We find that almost one in four teleost gene phylogenies in the Ensembl database (3,391) are inconsistent with their syntenic contexts. For 70% of these gene families (2,387), we were able to propose an improved phylogenetic tree consistent with both the molecular substitution distances and the local syntenic information. We show that these synteny-guided phylogenies are more congruent with the species tree, with sequence evolution and with expected expression conservation patterns than those produced by state-of-the-art methods. Finally, we show that synteny-guided gene trees emphasize contributions of WGD paralogs to evolutionary innovations in the teleost clade.

## Introduction

Whole genome duplications (WGDs) are dramatic evolutionary events that result in the doubling of a species entire genome. Several ancient WGDs have occurred in land plants, fungi and animals, in which they represent a major source of functional innovation with long-term impact on the evolution of species (Jaillon et al. 2004; Van de Peer et al. 2017). Genome doubling events have also been uncovered in non-model organisms in recent years (Kenny et al. 2016; Sollars et al. 2017), with more still to be discovered. While many gene duplicates produced by WGDs are eventually lost, some are retained and thought to provide raw material for evolution to repurpose into new functions (Ohno 1970; Lynch and Conery 2000). For example, 35% of human genes still have ancient duplicates from two WGD events ~550 million years ago, which diversified essential multigene families including the MHC (immunity) or the four HOX clusters (anteroposterior development) (Sacerdot et al. 2018; Singh and Isambert 2019). Investigating the fates of duplicate genes in descendant species is crucial to understand how WGDs contribute to phenotypic robustness, adaptation and evolvability. This however requires the accurate identification of orthologs (descended from the same post-WGD gene copy) and paralogs (descended from different post-WGD gene copies) across species. WGD paralogs, also called ohnologs, must also be distinguished from paralogs that arose by small-scale duplications or retrotransposition (Hahn 2009).

Orthology and paralogy relationships between genes are generally inferred from a phylogenetic tree. This gene tree represents the most likely evolutionary history from a common ancestral gene based on existing gene sequences. Reconciliation with the species phylogeny then labels gene duplication and speciation events. These phylogeny-reconciled gene trees allow rigorous, model-based tests to examine the evolution of gene sequences and functions. However, gene trees are known to contain errors related to methodological, technical and biological factors (Som 2015). A prominent source of errors is low phylogenetic signal in sequences, i.e. insufficient numbers of substitutions to confidently support one gene tree topology over others (Rasmussen and Kellis 2007). To address this issue, species-tree-aware methods use proximity to the structure of the species tree, known as the reconciliation cost, to select among statistically equivalent gene tree topologies (Durand et al. 2006; Vilella et al. 2009; Rasmussen and Kellis, 2011; Szöllősi et al. 2013; Wu et al. 2013; Scornavacca et al. 2015). Because of computational trade-offs, these methods either rely on heuristic exploration of the gene-tree solution space and result in suboptimal trees, or have an intensive computational cost and are not applicable to large datasets. Critically, these limitations are enhanced in the presence of ancient whole genome duplications. Species descended from a WGD frequently have two paralogs or more per gene family, directly increasing the size of the solution space. Further, reconciliation methods do not account for the acceleration of gene loss rates shortly after the WGD, or for rate heterogeneity across species (Zwaenepoel and Van de Peer 2019).

However, we argue that WGDs also possess underexploited characteristics that can be leveraged to improve gene tree modelling. First, the known timing of well-supported WGD(s) can be integrated as prior knowledge to select or constrain gene tree topologies. Second, WGDs result in specific patterns of genome organisation, where pairs of sister regions share duplicated genes in conserved order throughout the genome. These double-conserved syntenic (DCS) regions can be readily uncovered by comparison with unduplicated outgroup genomes (Kellis et al. 2004; Jaillon et al. 2004). Because reconciled gene trees are unreliable, DCS patterns are commonly used to identify WGD duplicated genes with confidence (Byrne and Wolfe 2005; Catchen et al. 2009; Muffato et al. 2010). In multi-species studies, this evidence has typically been used to exclude gene families where synteny disagrees with the precomputed tree structure (Kassahn et al. 2009; Berthelot et al. 2014; Braasch et al. 2016). However, synteny has never been used to systematically select amongst alternative gene tree topologies in the context of a known WGD event. To the best of our knowledge, only two gene tree building methods are able to leverage evidence from gene organisation to correct orthology and paralogy relationships: SYNERGY, a Neighbor-Joining iterative tree building algorithm that is no longer available (Wapinski et al. 2007), and ParalogyCorrector, which identifies and corrects gene trees inconsistent with synteny information (Lafond et al. 2013). However, ParalogyCorrector is designed to remove unsupported duplication nodes and recover all true orthologs at the expense of paralogs, which makes it unsuited to WGD studies. Here, we present SCORPiOs (Synteny-guided <u>COR</u>rection of <u>P</u>aralogies and <u>O</u>rthologies), a synteny-guided gene tree building algorithm for WGD studies. SCORPiOs builds optimized, species-tree-aware gene trees consistent with known WGD events, local synteny context, and gene sequence evolution. We apply SCORPiOs to the teleost-specific genome duplication, dated 320 Mya (Jaillon et al. 2004) and find that almost one in four WGD-descended gene trees are incorrect. We propose a corrected, synteny-consistent tree for 70% of these gene families (2,387 of 3,391 synteny-inconsistent trees). Then, we show that the corrected trees emphasize how duplicate gene retention after the WGD has shaped developmental and signalling pathways involved in known phenotypic innovations in the teleost lineage.

## Results

### Synteny is informative to describe gene phylogenetic relationships after WGD

After a whole genome duplication, gene positions along chromosomes display a particular conservation signature (double-conserved synteny) that can be leveraged to identify and differentiate orthologous and paralogous duplicated regions across species (Kellis et al. 2004; Jaillon et al. 2004). This genomic organisation reveals gene trees that are inconsistent with their local syntenic neighbourhood (Figure 1A). To test whether this identification can be extended systematically to the entire genome, we asked whether local gene neighbourhoods significantly differ between orthologous and paralogous WGD gene copies in the absence of any correction. In theory, paralogous regions diverge immediately after the WGD event, in particular through massive gene losses, and should be more different in terms of gene retention, loss, and molecular evolution across species compared to orthologous genomic segments, which diverge at speciation (Figure 1B; (Scannell et al. 2006)). We compared the local genomic context of 2,394 duplicated gene pairs in zebrafish and medaka from unambiguous sequence-based gene trees (Methods) and found that orthologs indeed share more orthologous syntenic neighbours and have more similar local gene retention and loss patterns than paralogs, as expected (Figure 1C, Wilcoxon–Mann–Whitney tests, p < 2.2e-16). Local syntenic context is therefore informative to distinguish orthologs from paralogs, and can potentially be leveraged to systematically correct erroneous gene trees after a WGD event.

**Figure 1:**
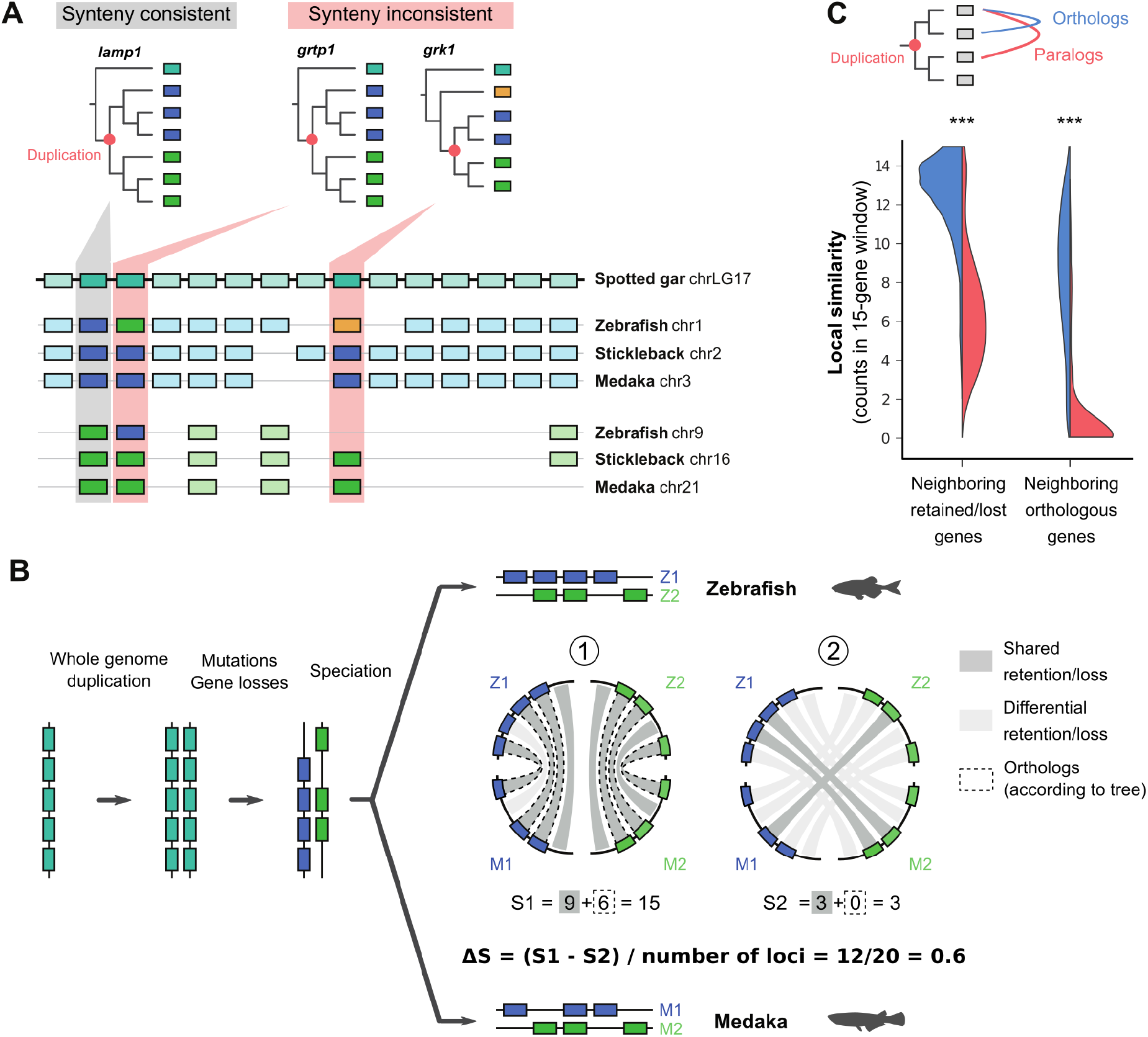
Local synteny context is informative to sort out gene duplicates after WGD. **A.** Gene trees and synteny context of the *lamp1*, *grtp1* and *grk1* gene families in teleosts and their non-duplicated outgroup (spotted gar). Gene trees are labelled with duplication nodes in red. Colors represent teleost genes annotated as orthologs in the original gene trees from Ensembl (blue/green, with yellow genes annotated as non-duplicated). The synteny context identifies that in some gene trees, zebrafish homologs are assigned to the wrong orthology groups (highlighted in red). **B.** Schematic evolution of a genomic segment after WGD, with gene colors as in A. After WGD, the duplicated segments evolve independently and accumulate mutations and gene deletions. Zebrafish and medaka genomic segments are compared under two scenarios: Z1/M1 and Z2/M2 (scenario 1), which corresponds to true orthologs, and Z1/M2 and Z2/M1 (scenario 2), corresponding to paralogs. Under each scenario, genes similarly retained or lost across species are represented by grey links, and genes annotated as orthologs are linked by dotted lines. True orthologous segments (1) share more similar patterns of gene retentions/losses and orthologies than paralogous segments (2), resulting in a high four-way similarity score ΔS. **C**. Local syntenic context can differentiate orthologs from paralogs. Distributions of the number of shared gene retentions/losses and annotated orthologs in a 15-gene window around zebrafish/medaka orthologs (in blue) and paralogs (in red), based on the original gene trees from Ensembl. Wilcoxon–Mann–Whitney tests, n=2,394, *** p < 0.001.

### SCORPiOs: Synteny-guided CORrection of Paralogies and Orthologies in gene trees

Integrating sequence and synteny information to build gene phylogenies is challenging, and current methods are not designed to deal with WGDs. To address this issue, we have developed SCORPiOs, an algorithm that improves gene trees in the presence of one or several WGD events, using information from synteny conservation patterns. SCORPiOs is coded in Python 3, implemented as a snakemake workflow (Köster and Rahmann 2012), and takes as inputs: (1) the gene positions in a set of paleopolyploid genomes and one or several unduplicated outgroups, (2) the species phylogeny with the putative WGD position(s), (3) the set of phylogenetic gene trees to be tested and amended for WGD consistency and (4) the corresponding gene sequence alignments. For convenience, SCORPiOs also includes an implementation of TreeBeST (Vilella et al. 2009) and can initialise the process from sequence alignments if precomputed gene trees are not available. The major steps of SCORPiOs are outlined in Figure 2 (see Supplementary Figure S1 for a detailed flowchart):

i. the identification of all potential duplicated genes and regions in each duplicated genome;
ii. the identification of orthologous segments between pairs of species, and by extension orthologous genes, by scoring local synteny similarity;
iii. for every gene family, the integration of the predicted orthology relationships in an orthology graph, and derivation of communities of orthologous genes across species;
iv. the correction of gene trees that are inconsistent with these orthology and paralogy constraints.

**Figure 2:**
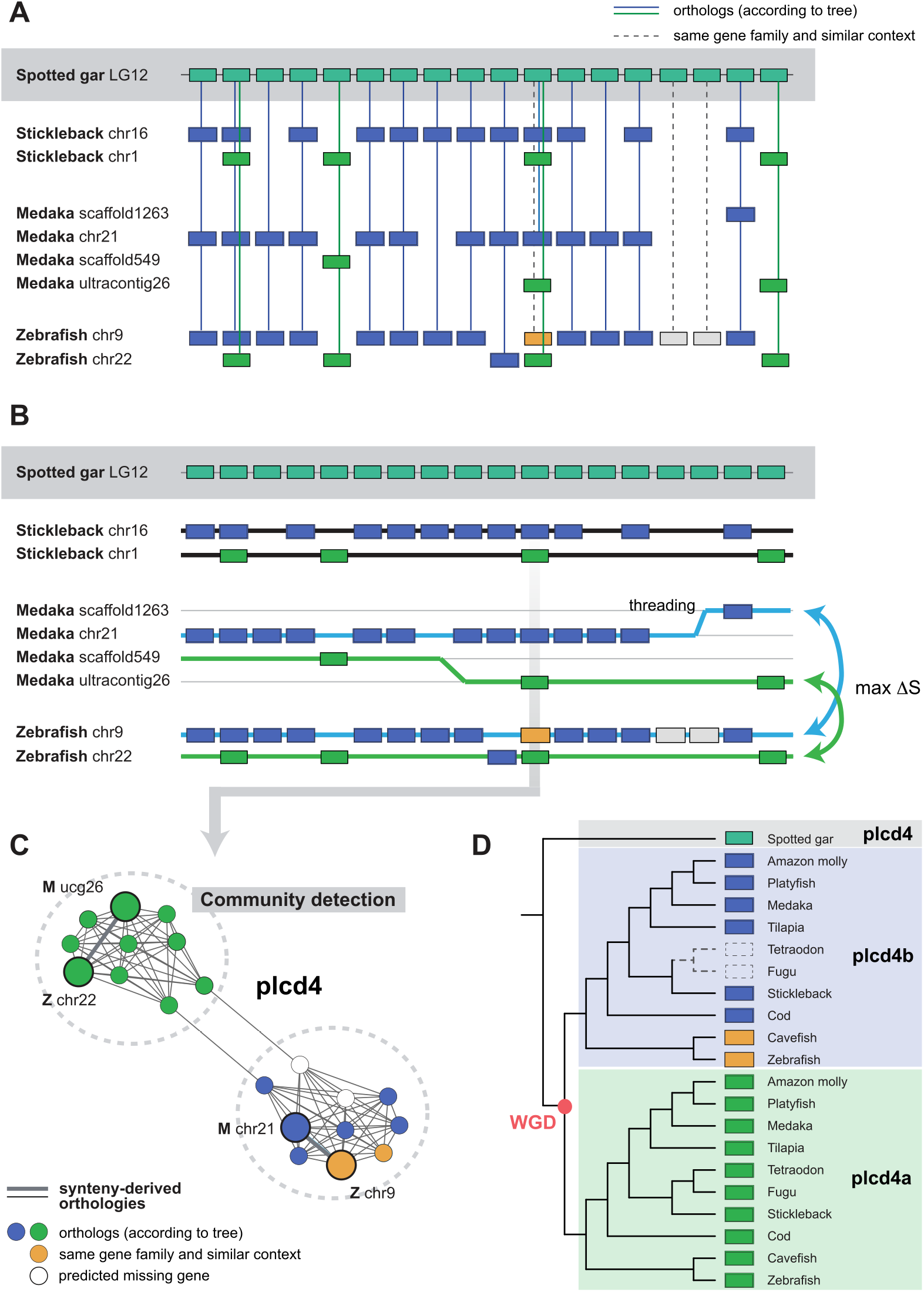
Overview of the SCORPiOs workflow. **A.** For every region in the reference non-duplicated genome, SCORPiOs identifies all potential orthologs in the duplicated genomes and builds a relaxed orthology table. Here, a section of the spotted gar linkage group 12, and genomic locations of its potential orthologous genes in three TGD teleosts (out of ten). For each gene family, color represents genes identified as orthologs in the original gene trees from Ensembl **B.** SCORPiOs maximizes the synteny context similarity ΔS to thread and pair orthologous genomic segments between duplicated species (see Supplementary Note 2). Green and blue threads show the highest-scoring configuration for the medaka/zebrafish comparison. **C.** For each individual gene, SCORPiOs integrates orthology links derived from the local synteny context across all species into a gene graph, where orthologous communities are identified. Here, the *plcd4* gene, whose genomic position is highlighted in B. Nodes represent homologs of the *plcd4* gene, colored as in A and B, where the medaka and zebrafish genes are highlighted as large circles. Links represent orthology relationships deduced from the similarity in syntenic contexts, with orthologies between zebrafish and medaka highlighted with a thicker line. SCORPiOs explicitly models nodes corresponding to missing homologs in the gene graphs (in white). **D.** If the original gene tree is not consistent with the synteny-derived orthologous gene communities, SCORPiOs proposes a corrected gene tree. Here, the corrected gene tree for the *plcd4* gene family, which includes zebrafish and cavefish homologs (in yellow) as part of the *plcd4b* subclade based on their genomic location (as seen in B).

The implementation details of these different steps in SCORPiOs are developed below.

### Comprehensive identification of homologous families after the WGD event

To identify duplicated regions within a genome, SCORPiOs takes advantage of patterns of ‘double-conserved synteny’ (DCS) with a non-duplicated outgroup. After the WGD, genomic regions of the non-duplicated outgroup give rise to two orthologous genomic segments in each duplicated ingroup species (Figure 1B). However, genomic rearrangements, gene losses or duplications, and errors in the initial gene trees can obscure these 2:1 orthologous relationships in modern genomes. Most WGD studies thus restrain themselves to genomic regions where the WGD event is evident either from the gene trees or from highly conserved DCS patterns (Scannell et al. 2006; Kassahn et al. 2009; Berthelot et al. 2014; Inoue et al. 2015; Braasch et al. 2016). As a consequence, significant subsets of gene families are de facto discarded from analysis (54% not considered in Inoue et al. 43% in Braasch et al.). SCORPiOs addresses this problem by first identifying all potentially WGD-descended gene duplicates. SCORPiOs uses a loose definition of orthology to build a comprehensive orthology table between the non-duplicated reference genome and all duplicated genomes from the initial gene trees (Figure 2A, Supplementary Note 1, Supplementary Figure S2). This table retains all gene copies since the ingroup/outgroup speciation node, as well as any other homologs from the same gene tree with a loosely similar syntenic context. Unlike other methods, SCORPiOs therefore first strives to define a comprehensive gene set from each gene family to be searched for potential orthology and paralogy relationships, instead of pruning all genes that do not belong to predefined duplicated segments.

### Identification of orthologous genomic segments between pairs of species

As a second step, SCORPiOs reconstructs ancestral WGD-duplicated regions, which may have become eroded by genome evolution processes. SCORPiOs uses the non-duplicated outgroup as a proxy for the ancestral gene order, and scans the sorted orthology table using a sliding window of user-defined length and threads the ancestral duplicated segments in each descendant species (Figure 2B; Supplementary Note 2). These duplicated genomic segments are then compared between pairs of duplicated species to match orthologous segments. SCORPiOs scores the syntenic similarity between segments using the number of genes annotated as orthologs in the initial gene trees, as well as the pattern of gene retention and losses, as described in Figure 1. SCORPiOs is flexible and does not require perfect gene order conservation or contiguity to thread genomic segments, instead attempting to maximise the local similarity between both duplicated species (Supplementary Note 2). This results in a four-way comparison of threaded, scored genomic segments for every pair of duplicated genomes and every n-sized window in the orthology table (Figure 2B).

SCORPiOs interprets the differentiation in four-way similarity scores (ΔS) as a confidence index that two genomic segments can be assigned as orthologs between the two duplicated species. This information is then reflected back to the genes within those segments to identify putative orthologs and paralogs using the window of highest ΔS that they belong to (Figure 1B, Supplementary Note 2, Supplementary Figure S3).

### Orthology and paralogy constraints across all species

As the previous step assigns orthologs and paralogs between pairs of species, SCORPiOs then integrates those results to define groups of orthologous genes across all species under study. SCORPiOs exhaustively performs all pairwise comparisons between duplicated genomes as described above, and constructs an orthology graph for every gene family (Figure 2C). SCORPiOs orthology graphs are unweighted graphs where two nodes (genes) are linked by an edge when they are inferred as orthologs based on their local synteny similarity. The graphs also include “placeholder” nodes, which correspond to gene copies that have been lost since the WGD but whose existence can be inferred from the synteny pattern (Supplementary Figure S4). In each graph, we then expect to find two similar-sized orthologous gene communities that are derived from the WGD. If synteny was perfectly informative and the process was error-free, we would expect these communities to be two independent, fully-connected cliques. In practice, we observe that some graphs do not result in two isolated communities due to inconsistencies in the orthology assignments. SCORPiOs then uses the Girvan-Newman algorithm (GN), a community detection algorithm that iteratively removes the most central edges of each graph to separate nodes in two communities (Girvan and Newman 2002). In cases where GN is unable to separate genes of a duplicated species in two communities, we apply the Kerningan-Lin algorithm (KL) (Kernighan and Lin 1970). KL is a heuristic aimed at finding two communities of similar sizes, that require cutting the fewest edges. If the KL solution separates the duplicated genes from the same species more frequently, it is preferred over the GN solution. At the end of this step, SCORPiOs has identified the two groups of orthologous genes that descend from the WGD, and consequently paralogous to each other, derived from local synteny information. Of note, one of these groups can be entirely made of placeholders, corresponding to the loss of one duplicate gene in all species (typically an early loss before the first speciation, Supplementary Figure S4).

### Gene tree correction and test

Next, SCORPiOs identifies and attempts to correct gene trees that do not fulfill the orthologies and paralogies from their synteny-derived orthology graph. First, SCORPiOs checks whether each gene tree contains a node from which the unduplicated outgroup gene diverges, followed by either (i) a node corresponding to the WGD, under which each subtree encompasses one of the two orthology communities from the orthology graph, or (ii) a single subtree, in cases where the same paralog is missing in all species (i.e. placeholder nodes subgraph). When this topological constraint is not verified in the original gene tree, SCORPiOs proposes a corrected, fully-resolved gene tree that is both species-tree and synteny-consistent. We explore the constrained topologies solution space using the fast distance-based approach implemented by the program ProfileNJ (Noutahi et al. 2016). Briefly, ProfileNJ independently resolves each multifurcated orthogroup, so as to minimize the reconciliation cost with the species-tree. SCORPiOs then compares the likelihoods of the original and corrected tree and accepts the correction if both trees are statistically equivalent (AU test (Shimodaira and Hasegawa 2001); branch lengths fitted with PhyML (Guindon et al. 2010); Methods). If ProfileNJ fails to find an adequate solution, SCORPiOs uses the TreeBeST-modified version of PhyML, which fits the gene sequences in each post-WGD subtree using maximum likelihood optimization while accounting for the species phylogeny topology (Vilella et al. 2009), and compares the corrected tree to the original one as above. We correct gene tree topologies only when the sequence similarity data is explained equally well (or better) by the synteny-aware tree: our correction approach is conservative, in the sense that it gives precedence to the likelihood of the molecular evolution model. Note also that SCORPiOs does not attempt to build a corrected tree for very large multigenic families (>1.5 genes per species in one or both post-WGD communities), as these are often over-aggregated families in input trees that SCORPiOs cannot solve. As a final step of the pipeline, SCORPiOs can re-graft the corrected subtrees into the original gene tree depicting the evolution of the whole gene family, and recompute branch lengths (Supplementary Note 3, Supplementary Figures S5-7). Additional available options when executing SCORPiOs are described in Supplementary Note 4.

### SCORPiOs corrects a significant fraction of Neopterygii subtrees in Ensembl

We applied SCORPiOs to gene trees from the Ensembl Compara database, version 89 (Vilella et al. 2009), which includes 70 vertebrate genomes, of which eleven are Neopterygii: ten duplicated teleost genomes, and the spotted gar as an outgroup to the teleost-specific genome duplication (TGD). This set represents a total of 21,431 gene trees reconciled with the vertebrate species tree. About 50% of genes are syntenic between spotted gar and teleost genomes, which makes it a suitable outgroup and proxy for the ancestral gene order (Methods).

We ran SCORPiOs in iterative mode, which corrects the gene trees a first time, thus improving the quality of the synteny information, and then again until convergence (window size: 15 genes, with a step of 1 gene). For this dataset, two iterations were sufficient to reach convergence (see Supplementary Tables S1-2 for detailed, by iteration, results). In brief, in iteration 1, we identified 15,476 Neopterygii loose orthology groups within the 21,431 gene trees. SCORPiOs produced 14,576 orthology graphs, of which 10,172 (70%) were already subdivided into two isolated orthology cliques. The vast majority of families not producing graphs (86 %) are small families (< 3 genes), for which we do not build graphs, as they would result in the same tree topology (Supplementary Table S1-2). For 3,391 gene trees (22%), the orthologies and paralogies relationships were in disagreement with the synteny graph. In total, at the end of the two iterations, we were able to find a synteny-aware and statistically equivalent gene tree for 2,387 of these genes families, thus correcting 70% of all synteny-inconsistent gene trees (15% of Ensembl Neopterygii subtrees). Interestingly, for 672 of the gene trees that we correct (28%), the new tree is better supported by the sequences and results in a significantly higher likelihood (AU test, α = 0.05). This may reflect difficulty of the Ensembl pipeline to explore the tree topology space due to the high number of species in the database, as previously described (Wu et al. 2013). Furthermore, we also applied SCORPiOs to version 94 of the Ensembl Compara database, containing 47 teleost genomes and a total of 43,491 gene trees. With this increase in phylogenetic coverage, a higher proportion of gene trees was inconsistent with synteny (7,168 synteny inconsistent gene trees for 15,760 gene families, 45%), and could be improved by SCORPiOs (3,681 corrected gene trees, 23%). Again, this result is in line with the idea that exploration of the topology space becomes limited for larger trees, inducing more errors. Altogether, we demonstrate that insight from synteny can improve a significant fraction of gene trees in the presence of many gene duplicates after a WGD event.

### Validation of SCORPiOs trees and comparison to existing methodologies

SCORPiOS corrects gene trees by providing synteny-derived orthology and paralogy constraints to existing tree building programs, ProfileNJ and TreeBeST PhyML. To evaluate how SCORPiOs improves over the state of the art, we compared the 2,387 corrected teleost gene trees to alternatives topologies obtained with other methodologies. First, we used RAxML (Stamatakis 2014) to build a pure maximum likelihood (ML) tree for each gene family, as a standard to compare against the likelihood of other trees. Second, we included the original, uncorrected trees from Ensembl Compara (Vilella et al. 2009), which are phylogeny-reconciled consensus trees build with the TreeBeST pipeline. Last, we tested ParalogyCorrector, the only other existing methodology that corrects gene trees using synteny information (Lafond et al. 2013). To compute ParalogyCorrector trees, we provided our synteny-derived orthologies constraints along with the original Ensembl subtree. ParalogyCorrector then finds a new gene tree minimizing the differences to the original tree while satisfying the orthology constraints (but not necessarily paralogy), while SCORPiOs fully recomputes each paralog subtree under the corrected duplication node.

We first evaluated whether each tree is well-supported by the sequence alignment (AU test, α= 0.05; Figure 3A). As expected, RAxML solutions are systematically the best fit to the sequence data. However, SCORPiOs is able to find a solution as good as the pure ML fit for 61% of gene trees, which is significantly better than the original Ensembl trees (47%; proportion test, p < 2.2e-16) or ParalogyCorrector (51%; proportion test, p < 2.2e-16). Additionally, the SCORPiOs tree is a better fit to the gene sequence information than either the Ensembl or the ParalogyCorrector trees for 9.5% of the 2,387 test gene families.

**Figure 3:**
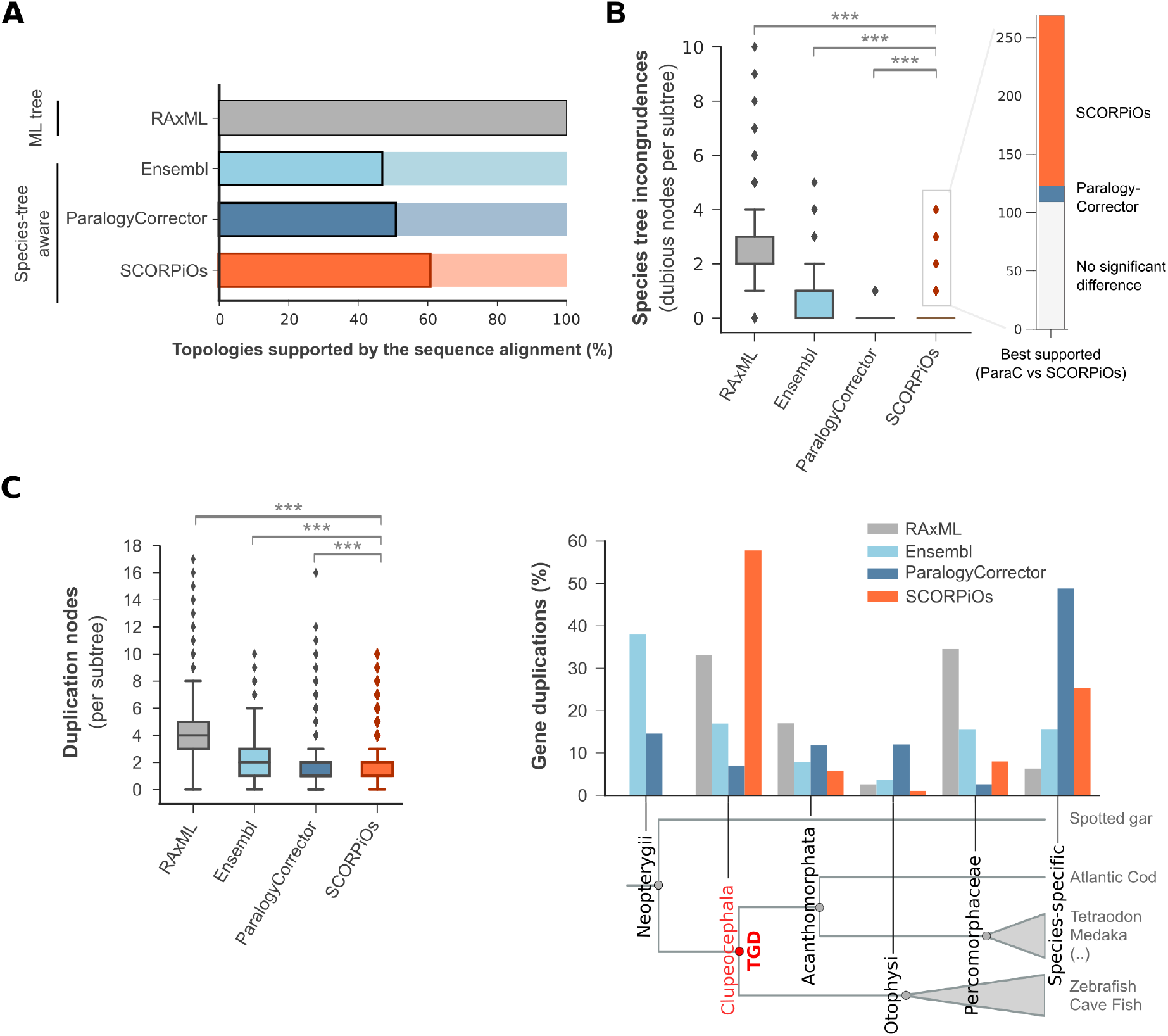
SCORPiOs performs better than state-of-the-art methods on WGD gene trees. **A.** SCORPiOs produces gene trees with higher phylogenetic support than either ParalogyCorrector or the Ensembl Compara pipeline. Bars represent the fraction of trees where the topology proposed by either of the three species-tree aware methods is not significantly less likely than the maximum-likelihood, non-reconciled solution from RAxML (AU-tests, p ≥ 0.05). **B.** Divergences from the species tree in gene trees produced by each method, assessed by the number of dubious duplication nodes per teleost subtree. For trees where SCORPiOs introduces more dubious duplication nodes than ParalogyCorrector, we found that the solution from SCORPiOs is typically better supported by molecular evolution (right panel). **C.** Number and position of gene duplications in the trees. Left: distribution of the number of duplication nodes for each method. Right: distribution of duplication nodes in the species tree for each method. The Clupeocephala ancestor corresponding to the expected TGD node is highlighted in red. Intermediate ancestors with fewer than 5% of duplications for all methods are not displayed.

We then assessed whether the gene trees are concordant with the known species phylogeny. For each software, we identified the species-tree inconsistent nodes, previously referred to as “dubious nodes” or “non-apparent duplication nodes” ((Vilella et al. 2009; Lafond et al. 2014); Methods). We find that SCORPiOs outperforms both RAxML and the original Ensembl trees in terms of species-tree congruence (Wilcoxon signed rank test, p-values < 2.2e-16, Figure 3B). Both SCORPiOs and ParalogyCorrector produce trees that are highly congruent to the species phylogeny, with a slight advantage to ParalogyCorrector (Wilcoxon signed rank test, p < 2.2e-16, 88.6% and 99.7% of fully-congruent trees, respectively; proportion test p < 2.2e-16). However, the non-congruent solution from SCORPiOs was better supported by sequence information for 146 of 269 gene families for which ParalogyCorrector reported a fully-congruent solution, suggesting that the additional duplication nodes inferred by SCORPiOs are largely correct.

Lastly, we assessed whether gene trees inferred by all four methods are parsimonious by calculating their total number of duplication nodes. We found that SCORPiOs infers the lowest number of duplications in gene trees, compared to all other methods (Wilcoxon signed rank test, p-values < 2.2e-16, Figure 3C). By design, SCORPiOs explicitly replaces the whole-genome duplication node at its known location in the tree when both gene copies survive in modern genomes (Figure 3D), resulting in fewer inferred duplications downstream of the WGD node. In contrast, ParalogyCorrector places duplications closer to the leaves and wrongly identifies most WGD duplicates as species-specific duplications. For 38% of Ensembl gene trees, the duplication is placed at the Neopterygii node and incorrectly encompasses the spotted gar gene (non-duplicated outgroup). In conclusion, SCORPiOs significantly improves gene trees in an evolutionary context where other methods struggle, due to the high number of gene copies and non-parsimonious tree structures created by WGD events.

### SCORPiOs correction improves correlation of orthologous gene expression

In addition to their consistency with sequence evolution and phylogeny, we evaluated whether orthologs corrected by SCORPiOs are functionally more similar than paralogs, as previously reported (Koonin 2005; Altenhoff et al. 2012; Chen and Zhang, 2012). We used gene expression data from 11 tissues in zebrafish and medaka (Pasquier et al. 2016) to investigate whether SCORPiOs-corrected zebrafish/medaka orthologs display higher similarity in expression patterns (Methods). For instance, SCORPiOs modified the orthology relationships in the cxcl12 gene family for zebrafish and medaka (Figure 4A), grouping together medaka cxcl12a and zebrafish cxcl12b in one orthology group and medaka cxcl12b and zebrafish cxcl12a in the other based on their local syntenic surroundings. Indeed, gene expression patterns support the orthology reassignment, with one gene copy expressed in bones, brain, embryo, gills and liver, and the other expressed at high level, predominantly in kidney, in both species.

**Figure 4:**
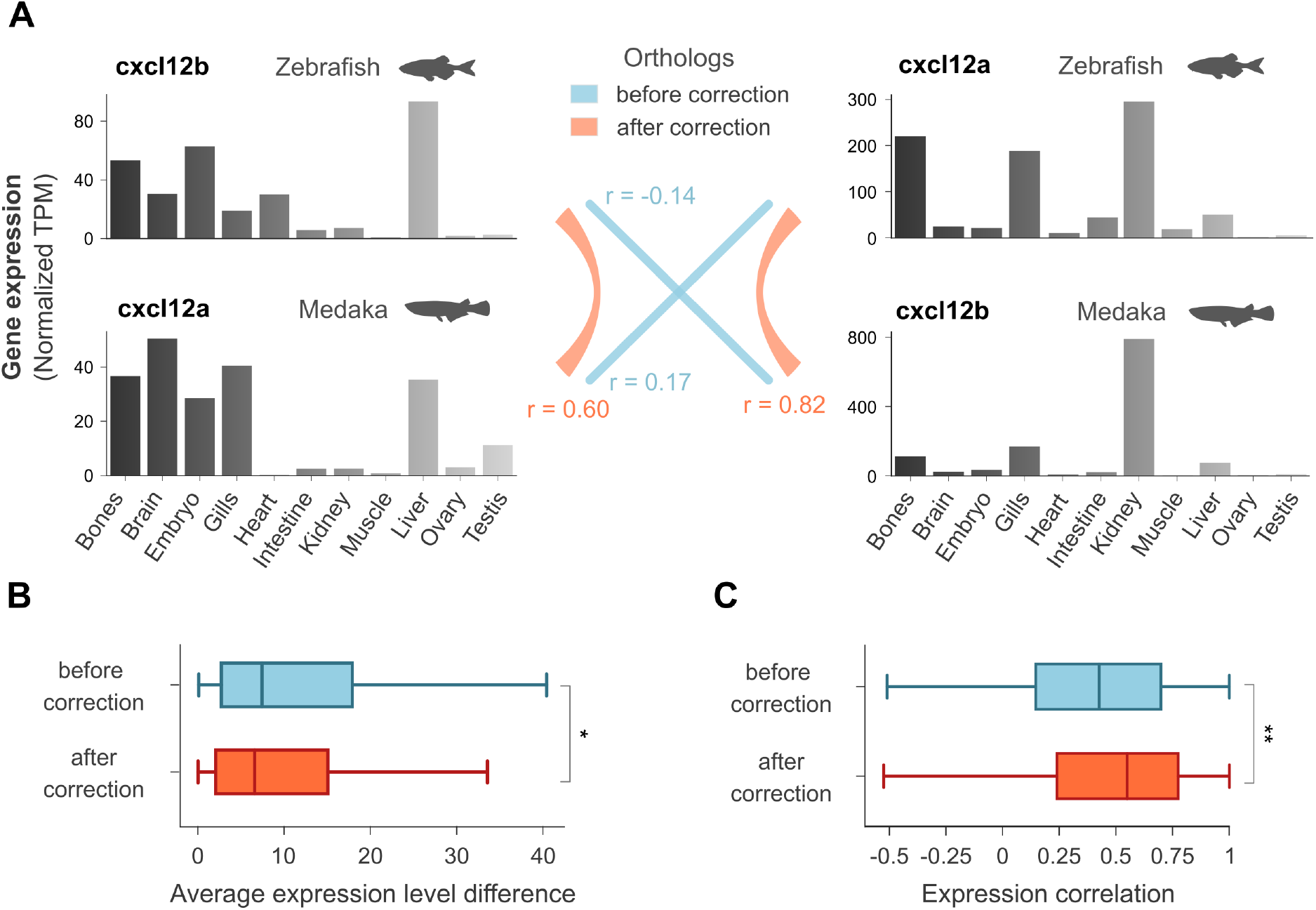
Functional similarity of orthologous genes after SCORPiOs correction. **A.** Expression of zebrafish and medaka *cxcl12* homologs in 11 tissues (quantile-normalized transcripts per million, TPM). Orthology relationships and expression level correlations before and after correction by SCORPiOs are noted in colour (r: Pearson correlation coefficient). **B.** Difference in average expression levels across 11 tissues between orthologous zebrafish/medaka genes, before and after correction (paired Wilcoxon test, n = 210, * p < 0.05). **C.** Average correlation of expression levels for orthologous zebrafish/medaka genes before and after correction (paired Wilcoxon test, n = 210, ** p < 0.01).

Overall, SCORPiOs modified the orthology relationships between zebrafish and medaka for 761 gene families. Of these, 210 correspond to orthology/paralogy reassignments (as in Figure 4A), while the remaining 551 correspond to removal of errors or addition of new homology relationships. The correction increased the number of 1-to-1 orthologs between zebrafish and medaka (13,463 vs. 14,008) as well as the total number of genes with an ortholog in the other species (16,150 zebrafish and 15,543 medaka genes with an ortholog before correction, vs. 16,316 and 15,748 after). For the 210 gene families where medaka and zebrafish orthologies were reassigned, we find that orthologs are expressed at closer average levels after correction (Wilcoxon signed rank test, p-value = 0.0171, Figure 4B) and also significantly more correlated across tissues (Wilcoxon signed rank test, p-value = 0.0050, Figure 4C). These results support that SCORPiOs measurably improves orthology and paralogy relationships. Additionally, they suggest that erroneous orthology relationships may obfuscate functional investigation of gene evolution after genome duplication, especially when their effects get compounded over dozens of species and thousands of gene families, as reported above for teleosts.

### SCORPiOs correction emphasizes WGD contributions to evolutionary innovations in teleost fish

Numerous studies have suggested a link between the function of a gene and retention or loss after a WGD. Yet, in the absence of systematic gene tree correction methods, the fate of TGD duplicates has only been investigated in a restricted set of ~6,000 high-confidence teleost gene families (Kassahn et al. 2009; Inoue et al. 2015; Braasch et al. 2016), potentially introducing biases in subsequent conclusions. Here we used the full set of 21,431 gene trees from Ensembl corrected by SCORPiOs to investigate gene retention across ten teleost species (zebrafish, cavefish, tetraodon, fugu, stickleback, medaka, tilapia, platyfish, amazon molly and cod). We classified genes into three categories with respect to their fate after the TGD (Methods). Briefly, we grouped genes retained in two copies across all 10 teleost species (“systematic ohnologs”, n = 1,828), genes found in single copy in all species (“singletons”, n = 13,895) and genes retained in two copies in at least one teleost species but not in all (“facultative ohnologs”, n = 7,265) (Supplementary Figure S8). We then used expression levels and functional annotations in zebrafish to explore how gene function relates to evolutionary trajectory after the TGD (Methods).

Overall, we find that singletons have slightly higher average expression levels and broader expression patterns than both systematic and facultative ohnologs (Wilcoxon–Mann–Whitney test, p < 0.001, Figure 5A, Supplementary Figure S9, Methods). This is in line with previous observations on paralogs and may reflect cases of subfunctionalization between the two groups of ohnologs, for which duplicated genes have partitioned the ancestral function, becoming expressed in fewer tissues and/or at lower level (Huminiecki and Wolfe 2004; De Smet et al. 2013; Guschanski et al. 2017). We next investigated whether tissue-specific singletons and ohnologs display preferential expression in different tissues, reflecting different contributions to teleost evolution. We find that tissue-specific systematic ohnologs are overrepresented in brain, heart and muscle, and depleted in liver, intestine, ovary and testis, compared to all zebrafish tissue-specific genes (tau > 0.9; hypergeometric tests, corrected p < 0.05, Figure 5B). In contrast, tissue-specific singletons genes are overrepresented in liver, kidney, intestine and testis (Figure 5B). For facultative ohnologs, we observe an enrichment in brain-specific genes, but also in muscle-specific expression (Figure 5B). Interestingly, enrichment in brain and heart specific genes, as well as depletion in liver and testis specific genes, have been already observed in human ohnologs retained after the 1R and 2R vertebrate WGDs (Guschanski et al. 2017). This result ties in with previous reports that some gene families and functional categories are recurrently amplified by independent WGDs, possibly because they offer adaptive advantages when duplicated en masse (van Hoek and Hogeweg 2009; De Smet and Van de Peer 2012).

**Figure 5:**
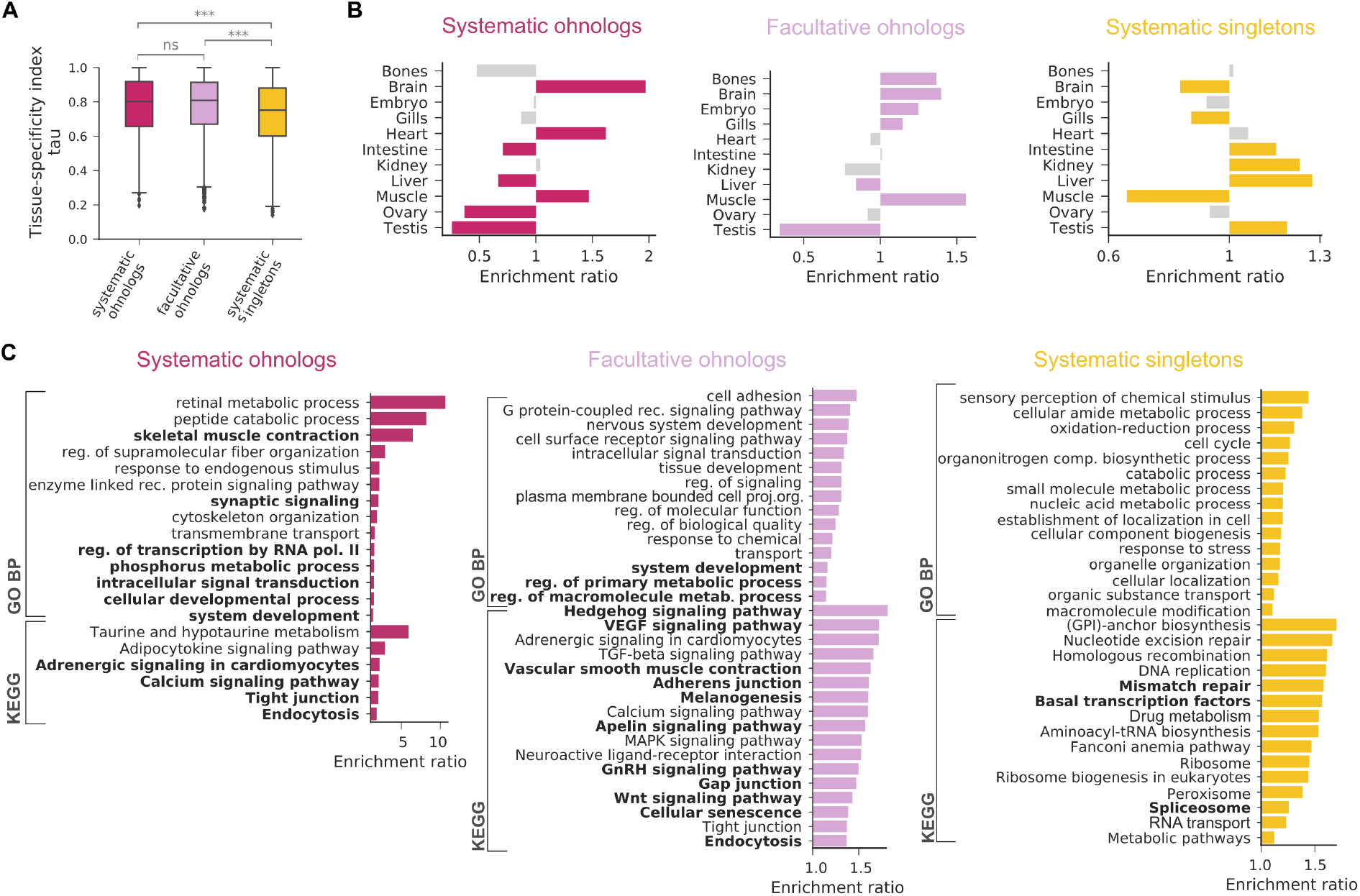
Functional analysis of genes with different evolutionary trajectories after the TGD. **A.** Tissue specificity of zebrafish genes with different evolutionary trajectories. Systematic ohnologs: WGD duplicates retained in two copies in all ten teleost species under study (n = 1,828). Facultative ohnologs: WGD duplicates retained in two copies in at least one species (n = 7,265). Systematic singletons: WGD duplicates returned to single-copy state in all ten species (n = 13,895). **B.** Preferential tissue of expression for tissue-specific genes (tau > 0.9) from each evolutionary trajectory. Colors denote statistical significance (hypergeometric test with BH correction, p < 0.05). **C.** Gene Ontology Biological Process and KEGG pathways enrichments for tissue-specific genes from each evolutionary trajectory (hypergeometric test with BH correction, p < 0.05). Bold: only enriched after gene tree correction with SCORPiOs.

Additionally, we investigated whether TGD ohnologs and singletons belong to different biological pathways using Gene Ontology Biological Processes (GO BP) and KEGG pathway enrichment analyses (Methods). Systematic and facultative ohnologs are enriched in general molecular processes previously found in WGD duplicates, linked to transcriptional regulation and metabolic processes, as well as terms related to the nervous system, consistent with the brain-specific expression patterns reported above (Figure 5C, Supplementary Tables S3-6) (Blomme et al. 2006; Inoue et al. 2015; Singh et al. 2015; Li et al. 2016; Pasquier et al. 2017) In contrast, singletons are enriched in housekeeping functions, with some of the most significant GO terms being “cell cycle”, “nucleic acid metabolic process” and “cellular localization”, along with the KEGG pathways “Ribosome” and “DNA replication” (Figure 5C, Supplementary Tables S7-8) (De Smet et al. 2013; Li et al. 2016). However, we discover here that systematic duplicates are also enriched in more specific functions, especially related to retina physiology. Interestingly, the teleost WGD coincides with functional innovations in the retina specific to this clade, where photoreceptor cells are organised in a regular pattern described as a ‘cone mosaic’ (Lyall 1957; Engström 1963; Sukeena et al. 2016). These results are mirrored in the KEGG analysis with the overrepresentation of the taurine and hypotaurine metabolism pathway, suggested to have a functional role in the teleost retina (Lima et al. 1998; Omura and Inagaki 2000). Our results therefore support that the amplification of retinal genes during the teleost WGD was important in the acquisition of this evolutionary innovation.

Finally, some functional enrichments become prominent only after gene tree correction with SCORPiOs (in bold on Figure 5). In particular, we observe an enrichment for both systematic and facultative ohnologs towards terms related to the circulatory system (“Adrenergic signaling in cardiomyocytes”, “Vascular smooth muscle contraction”, “VEGF signaling pathway”; Figure 5C, Supplementary Tables S4 and S6). This extends broader support to a previous report that TGD-derived duplicates, especially those of the *elastin* gene, have led to morphological sophistication of the teleost heart and circulatory system (Moriyama et al. 2016). Lastly, genes in the facultative ohnolog category are enriched in the “Melanogenesis” pathway, also consistent with the expansion of the pigmentation repertoire after the TGD (Lorin et al. 2018).

Taken together, our results suggest a strong contribution of TGD duplicates to functional novelty in this clade, mediated by the fixation of specialized duplicated genes. Importantly, many of these enrichments were fully obscured by errors in the gene evolutionary histories downloaded from as respected a reference database as Ensembl, which is widely sourced for comparative and evolutionary studies (Alföldi and Lindblad-Toh 2013; Herrero et al. 2016). SCORPiOs therefore fulfils its purpose in the arsenal of tree building tools and has potential to dramatically further investigations into the evolutionary and functional outcomes of WGD events.

## Discussion

Gene duplications have long been recognized as a major provider of raw material for molecular evolution and functional novelty (Ohno 1970; Lynch and Conery 2000). Whole-genome duplications have a substantially impact on genome evolution because they generate redundant copies for all genes, although only a fraction of these duplicates is retained over long evolutionary times. Gene retention and loss after a WGD is a poorly understood interplay between functional redundancy, increased evolvability and mutational cost. Several complementary models of gene evolution have been proposed to account for duplicate retention after WGD, including neofunctionalization, where one copy acquires a new function while the other maintains the original one (Ohno 1970; Lynch and Conery 2000); subfunctionalization, where both copies partition the ancestral function between themselves (Force et al. 1999); dosage balance, where subunits of macromolecular complexes are maintained as duplicates to ensure proper stoichiometry between interacting partners (Blomme et al. 2006; Veitia et al. 2008; Makino and McLysaght 2010); and cost of deleterious mutations, impeding pseudogenization and loss (Gout et al. 2010; Singh et al. 2012). Overall, the relative contributions of these processes to short- and long-term gene evolution remain unclear, although it is generally agreed that the evolutionary fate of genes following WGDs is tightly intertwined with their ancestral functions.

Polyploidisations are widespread through eukaryotic evolution, representing many independent opportunities to characterize gene evolution after WGD. Yet, WGDs represent a serious challenge to current gene tree reconstruction methods due to the high volume of duplicates and gene losses that they produce. As a result, incertitudes in gene phylogenies have been a limiting factor to all WGDs studies, whether they investigate the incidence and timing of ancient WGDs (Van de Peer et al. 2010; Ruprecht et al. 2017; Zwaenepoel and Van de Peer 2019), biases in duplicate gene retention (Scannell et al. 2006; Kassahn et al. 2009; Inoue et al. 2015), or genome organisation evolution (Varadharajan et al. 2018). Studying WGD occurrences and consequences calls for the development of specific methodologies to characterize the complex histories of WGD genes. Illustrating on-going efforts, the recently published WHALE approach accounts for incertitude in gene tree reconciliations when identifying plausible WGDs in a species tree (Zwaenepoel and Van de Peer 2019). SCORPiOs fills another methodological gap by integrating insights from genome evolution to improve reconciled gene trees.

Genome evolution operates through three major mechanisms: nucleotide substitutions, gene duplications and losses, and genomic rearrangements. To date, integrating these different evolutionary events into a unified framework remains an open challenge (Chauve et al. 2013). The inference of reconciled gene trees, which jointly depict the history of substitutions and gene gains and losses, has been growingly addressed in recent years (Szöllősi et al. 2015). Synteny conservation has the potential to neatly complement sequence similarity in gene evolution studies, because gene order evolves via independent mechanisms, and is more resilient to saturation at deep evolutionary times (Rokas and Holland 2000). Genome organisation information still remains difficult to incorporate in gene phylogenies, largely due to the lack of well-supported evolutionary models (Chauve et al. 2013) and the need for contiguous genome assemblies. The most notable effort to use extant synteny to correct gene trees in a general context showed mixed results (ParalogyCorrector within the RefineTree framework (Lafond et al. 2013; Noutahi et al. 2016)). However, we show here that in contexts where additional priors on genome organisation can be leveraged, synteny patterns can be highly informative and effectively improve reconciled gene trees. As high-quality reference genome assemblies are becoming affordable and straightforward, we expect that synteny will become increasingly useful to gene history resolution in an ever more complex comparative genomics landscape. For instance, synteny-aware methods will allow the investigation of other significant biological events, such as gene conversions, which introduce discordances in the history of a gene sequence and the history of its locus.

Lastly, assessing the quality of gene trees remains a challenging task, simply because the true evolutionary history of a gene is unknown. While statistical likelihood is widely used as a goodness-of-fit criteria to evaluate gene trees, the tree of maximum likelihood according to sequence evolution is frequently incorrect (Shimodaira 2002; Szöllősi et al. 2015). It is generally assumed that the correct tree falls within an interval of equally supported trees, but numerous factors can invalidate this hypothesis, ranging from errors in sequences or their alignment to unrealistic assumptions of evolutionary models. Consequently, other goodness-of-fit metrics have been introduced, including measures of species-gene tree discordance, parsimony of the duplication and loss scenario, distances to gold standard or simulated trees, functional similarity of orthologs, and power to reconstruct ancestral genomes (Altenhoff et al. 2016; Noutahi et al. 2016). Their use has been heterogeneous across studies, guided by specific aims, relevance and feasibility. Here, we validate SCORPiOs on real data, taking full advantage of computable metrics to demonstrate the improved quality of SCORPiOs corrected trees. In the future, efforts towards standardized benchmarking, as led by the Quest for Orthologs community, will be instrumental in producing ever more accurate gene phylogenetic trees.

## Methods

### Synteny similarity of WGD orthologs and paralogs in the absence of tree correction

Gene trees constructed with TreeBeST were downloaded from Ensembl v.89 (Vilella et al. 2009). We extracted a set of 2,394 high-confidence, WGD-descended homologous gene pairs in zebrafish and medaka using synteny criteria similar to approaches in (Kassahn et al. 2009; Braasch et al. 2016). Well-defined WGD duplicated regions were identified in the medaka and zebrafish genomes, where one contiguous 15-gene window in the spotted gar genome has homologs on exactly two different chromosomes in both medaka and zebrafish. Medaka and zebrafish gene pairs were considered high-confidence WGD duplicates when they are located at the midpoint of one of these 15-gene windows. The orthology and paralogy relationships between those gene copies were extracted from the original gene trees (1696 orthologous and 698 paralogous gene pairs). For each medaka-zebrafish gene pair (orthologs or paralogs), we counted across the 15-gene window: (i) the number of orthologs genes between medaka and zebrafish, according to the original trees, and (ii) the number of homologs to spotted gar genes similarly retained or lost in both species.

Genome-wide synteny conservation was calculated between spotted gar (used as the outgroup) and other teleost genomes in the absence of tree correction using PhylDiag with default parameters (Lucas et al. 2014).

### Gene tree comparisons

Nucleotide sequence alignments containing teleost fish sequences were downloaded from Ensembl v.89 and pruned of non-Neopterygii sequences. For each tree topology inferred by either TreeBeST, ParalogyCorrector or SCORPiOs, phylogenetic likelihood was computed with PhyML using the HKY85 model (Guindon et al. 2010). Likelihoods for alternative topologies were compared using the Approximately Unbiased (AU) test implemented in Consel, at *α* = 0.05 (Shimodaira and Hasegawa 2001).

Dubious nodes (or non-apparent duplication nodes) are gene duplications inferred by the reconciliation procedure where no descendant species actually contains two genes copies. They correspond to inconsistencies between the gene and species trees and are likely errors in the gene tree topology. To find dubious nodes, we used treebest sdi to reconcile gene trees with the species tree and identified all duplication nodes with a confidence score of 0 (Vilella et al. 2009).

### Gene expression analysis

We used RNA-seq datasets from the PhyloFish database (Pasquier et al. 2016), which provides transcriptomes in fish for the following tissues: bones, brain, embryo, gills, heart, intestine, kidney, liver, muscle, ovary and testis. We used kallisto (Bray et al. 2016) with default parameters to quantify transcript abundances for the full set of Ensembl transcripts in zebrafish and medaka. We summed Transcripts per Million (TPM) values of alternative transcripts to obtain a quantification of the expression of their corresponding gene. Finally, TPM values were quantile normalized within each species to obtain equivalent distributions of gene expression levels across tissues (Bolstad et al. 2003).

### Functional similarity of orthologs

We assessed the functional similarity of zebrafish-medaka orthologs before and after gene tree correction. From the 2,387 corrected gene families, we selected the subset of 210 trees where zebrafish and medaka orthologies were re-assigned by SCORPiOs. Orthologous gene expression levels were compared before and after correction using Pearson correlation and differences in mean expression across tissues. Average correlation and average expression difference before and after correction were compared using Wilcoxon signed rank tests. All tests are paired to ensure that results are unbiased by heterogeneous evolutionary rates across gene families.

### Evolutionary categories of genes

We used the corrected gene tree forest to classify zebrafish genes with respect to their evolutionary fate across species after the TGD. We extracted all teleost gene clades from the trees and used the duplication status of the root node to determine the fate of the descending genes across species. If the root node is not a duplication, then all genes returned to a single copy state after the TGD and we classify descending zebrafish genes as ‘systematic singletons’. If the root node is a duplication (corresponding to the TGD), and all descending species retained both duplicated copies (duplication confidence score = 1), we defined them as ‘systematic ohnologs’. Finally, if more than one but not all species retained the two copies (duplication confidence score < 1), we classify genes as ‘facultative ohnologs’. We excluded from this classification 2,086 zebrafish genes with no other teleost homologue in their respective subtrees.

### Expression of genes with different trajectories after the TGD

We used the tau index (Yanai et al. 2005) to assess the degree of tissue specificity of the expression of zebrafish genes. Tau varies between 0 and 1, where 0 means broad expression and 1 specific expression. We defined tissue-specific genes as genes with a tau index > 0.9 and tested for enrichment of genes specific to particular tissues using hypergeometric tests with Benjamini & Hochberg correction for multiple testing.

### Functional enrichment of genes with different trajectories after the TGD

We used WebGestalt (Liao et al. 2019) to search for functional enrichment in each evolutionary category of genes, with all zebrafish protein coding genes as background. For systematic ohnologs, 786 and 496 out of 1,828 genes were mapped to an annotation in GO BP and KEGG pathway respectively, 5,496 and 3,392 out of 13,895 in systematic singletons, and 2,698 and 1,625 out of 7,265 in facultative ohnologs. WebGestalt uses hypergeometric tests, corrected for multiple testing with the Benjamini & Hochberg procedure, to test for significant enrichment. We report significant enrichments at a threshold of corrected p-value < 0.01. For visualisation purposes (Figure 5C), we reduced redundancy of functional GO terms to a maximum of 15, using the weighted set cover method implemented in Webgestalt. In all cases, the reduced set covers more than 97% of the total genes in each category. We repeated the analysis starting from the uncorrected Ensembl gene forest to determine if the enriched annotations differ after applying SCORPiOs.

## Availability and implementation

SCORPiOs is coded in Python 3 and implemented as a snakemake workflow, supported on Linux and macOS. Code is publicly available on Github at https://github.com/DyogenIBENS/SCORPIOS. SCORPiOs is distributed under the GNU GPLv3 license.

## Acknowledgements

We thank Pierre Vincens for the coordination of computing resources and all members of the GenoFish consortium for fruitful discussions.

