## Supplementary Material for "Synteny-guided resolution of gene trees clarifies the functional impact of whole genome duplications"

**Supplementary Notes**

Supplementary Note 1: Construction of the Orthology Table
Supplementary Note 2: Threading ancestral duplicated segments
Supplementary Note 3: Regrafting corrected subtrees into complete gene trees
Supplementary Note 4: SCORPiOs additional options: Iterative mode, multiple outgroups, multiple WGDs and more

**Supplementary Tables**

Supplementary Table S1-2: SCORPiOs correction details on Ensembl89
Supplementary Tables S3-8: GO and KEGG pathway enrichment for genes with different evolutionary trajectories after the TGD

**Supplementary Figures**

Supplementary Figure S1: SCORPiOs workflow
Supplementary Figure S2: Orthology between duplicated species and the outgroup on the basis of synteny conservation
Supplementary Figure S3: Example for the computation of a ∆S score between threaded segments of two duplicated species
Supplementary Figure S4: Example of a SCORPiOs tree for a gene family with singletons in all species
Supplementary Figures S5-7: Examples of SCORPiOs subtrees regrafting
Supplementary Figure S8: Evolutionary categorization of zebrafish genes
Supplementary Figure S9: Average expression of zebrafish genes with different evolutionary trajectories after the TGD

**Supplementary Note 1: Construction of the Relaxed Orthology Table**

We detail here SCORPIOs' two-step procedure to exhaustively sample potential orthologous genes between a non-duplicated species (outgroup) and a group of duplicated species (ingroups).

The first step consists in identifying orthologies supported by phylogenetic sequence evolution in the original gene trees, i.e. homologs that split at the ingroup/outgroup speciation node. In practice, for each clade of ingroup genes in a gene tree, we search for an outgroup gene that effectively split from it through a speciation or a dubious duplication node (p.i). If several outgroup genes match criterion (p.i), in a many-to-1 (or many-to-many) relationship, we arbitrarily select the outgroup gene closest to the duplicated species genes in the tree, in terms of topological then branch-length distance. If no outgroup genes match criterion (p.i), we search if the tree contains a clade made only of ingroup and outgroup genes (p.ii). Again, we select one amongst all genes matched in (p.ii) based on distance in the tree.

Secondly, if (p.i) and (p.ii) are unsuccessful in uncovering an outgroup ortholog for ingroup genes, we search for synteny-supported orthologies, using neighbouring phylogenetic orthologies extracted in (p.i) and (p.ii). In the gene tree, we search for the homologous outgroup gene sharing the highest number of syntenic neighbours with ingroup genes. If this synteny conservation is supported by at least 2 syntenic neighbours in duplicated species on average (based on orthologies from step p(i) and p(ii)) in a 30-genes local neighbourhood (s.i), we include the homolog in the orthology table. This threshold recovers most phylogenetically missed orthologies with a low false positive rate, based on a training set of high confidence orthologies and paralogies (Supplementary Figure 2). In case of ties in (s.i), we do not choose one outgroup gene, unless they are in-paralogs (young duplicates, i.e after ingroup/outgroup speciation). Otherwise, they are ancient duplicates in the outgroup, where we cannot identify the true orthologs amongst the two (or more) based on synteny conservation.

**Supplementary Note 2: Threading ancestral duplicated segments**

We detail here how SCORPiOs threads duplicated segments to reconstruct putative ancestral WGD-duplicated regions, for each window in the orthology table.

In an ideal situation, each genomic region in the unduplicated reference outgroup corresponds to exactly two genomic regions in duplicated species. In practice, genomic rearrangements, both in the outgroup and in duplicated species, have often eroded this perfect double-conserved synteny signature. For this reason, SCORPiOs starts by threading genomic segments to reconstruct the two paralogous ancestral regions in each duplicated genome. In every duplicated genome, we collect all segments sharing putative orthologs with the reference region in the outgroup, without any constraint on gene order conservation. If two gene homologs are found on the same chromosome but more than 100 genes apart, these regions are considered as two different segments. SCORPiOs ranks segments based on their number of orthologs with the outgroup, thus revealing the two major duplicated segments. We then investigate whether the remaining minor segments result from evolutionary fragmentation of the ancestral regions, and if so, which major segment they should be threaded with (Figure 2B). We test every possible configuration that does not contradict the following: (i) two segments with paralogous genes cannot be threaded into a single post-WGD region, and (ii) a maximum of two paralogs per gene family (ohnologs) are authorized in the resulting pair of threaded segments. As a consequence, we ignore any minor segments with additional homologs, which are discarded from the synteny analysis and will be grouped with their closest orthologs in the gene trees based on sequence evolution. Finally, all possible threading configurations are compared for any pair of duplicated species. We then select the combination that yields the most parsimonious scenario, where orthologous segments pairs are the most similar while paralogs differ, based on the synteny similarity ΔS score:

$$\Delta S = \frac{|Similarity(scenario1) - Similarity(scenario2)|}{NormalizationFactor}$$

with:

$$Similarity(scenario) = \sum_{pair \in [pair1, pair2]} PatternSimilarity\,(pair) + OrthologousNeighbours(pair)$$

and :

$$NormalizationFactor = 4 * SlidingWindowSize$$

An example for the computation of a ΔS score between threaded segments of two duplicated species is detailed in Figure S3. In effect, SCORPiOs uses each duplicated genome as a guide to select the most parsimonious threading scenario in the other species, under the assumption that many rearrangements or assembly fragmentations are independent between the two genomes. Comparisons of the same genome with different species can result in different optimal threading scenarios, which makes the process robust to threading errors. In this composite synteny similarity score, both measures (counts of retention/loss patterns and orthologous neighbours) are weighted equally. As they are both in the same scale (in [0, 1]), we found no advantage to using z-scores ($R^2$ >0.98). Retaining $\Delta S$ as a simple summing of counts is more computationally efficient as it is directly computable (versus having to store all counts values before z-score normalization).

**Supplementary Note 3: Regrafting corrected subtrees into complete gene trees**

As a final step of the pipeline, SCORPiOs can regraft the corrected sub-trees, which contain only genes from duplicated species and the reference outgroup, into the complete gene tree of the gene family, which can contain genes from other species as well as other gene sub-families. Within the SCORPiOs framework, corrected subtrees depict the evolutionary history of a single gene ancestrally duplicated through the WGD event, and its un-duplicated outgroup ortholog. The aim of the regrafting step is to effectively reinsert these corrected subtrees as a clade of genes into the complete gene trees, while rearranging other branches as little as possible. SCORPiOs then recomputes branch lengths for the complete gene tree as in the TreeBeST pipeline, using nucleotide CDS back-translated protein alignments with the Hasegawa-Kishino-Yano (HKY) model.

We distinguish two cases when regrafting subtrees:
(i) in the original tree, genes in the corrected sub-tree form a complete clade,
(ii) in the original tree, genes in the corrected sub-tree do not form a complete clade.

In case (i), regrafting is straightforward and simply consists in removing the original sub-tree and grafting the corrected version in its place (Supplementary figure 7). In case (ii), regrafting the subtree requires rearranging the topology of interleaved genes. Interleaved genes can either belong to species related to the reference outgroup, which should be included in the clade, or they can belong to other more distant outgroups or other gene sub-families, which should not be included in the clade. SCORPiOs distinguishes between these two situations to handle regrafting, using the species tree as a guide. First, it identifies genes to include in the corrected subtree clade, belonging to species either related to the reference outgroup or that branch between the reference outgroup and WGD species (Supplementary figure 8). These will remain unchanged in topology and the direct outgroup of the duplicated species corrected topology. Then, all other interleaved gene are pruned from the tree, keeping their topology relative to genes in the corrected tree unchanged and placed as outgroup of the corrected subtree clade (Supplementary figure 9). Note that SCORPiOs does not systematically check that these rearrangements do not result in a drop in tree likelihood, but such cases are rare in our experience (14% with significantly lower likelihood, 49% with significant likelihood improvement, 37% with no significant likelihood difference, based on the 932 Ensembl v.89 corrected trees with interleaved gene rearrangements). SCORPiOs aims at specifically improving WGD-duplicated genes phylogenies, and will not resolve other potential errors in trees. As such, SCORPiOs corrected gene trees should be considered as a gene tree forest with improved duplicated species subtrees.

**Supplementary Note 4: SCORPiOs additional options: Iterative mode, multiple outgroups, multiple WGDs and more**

We detail below several options that are available when executing SCORPiOs.

**Iterative correction.** SCORPiOs can run in iterative mode, meaning that SCORPiOs improves gene trees a first time, and then uses the corrected forest as input for a new correction run. Correcting gene trees improves orthologies accuracy, which in turn makes synteny conservation patterns more informative, improving its contribution to gene tree reconstruction. Usually, a small number of iterations (2-3) suffice to reach convergence. In order to reduce the computation time when executing a new correction run, SCORPiOs restricts itself to regions with updated synteny information – i.e. at iteration n + 1, only gene trees in the same window as a corrected tree in iteration n will be considered for correction.

**Multiple outgroup integration.** SCORPiOs can use more than one reference outgroup to correct gene trees for a given whole-genome duplication. Any non-duplicated species can be used as outgroup, but phylogenetically close outgroups should be preferred as synteny with duplicated species will typically be more conserved. Using more than one reference outgroup can be useful in two ways. First, gene families for which an outgroup homolog is missing will either (i) not be corrected by SCORPiOs, which requires an outgroup gene, or (ii) be corrected using a more distant, unrelated outgroup homolog in the gene family, which may cause errors. Using additional outgroups therefore improves the exhaustivity and reliability of the gene forest correction. Secondly, in genomic regions that have been perturbed by chromosomal rearrangements in the outgroup, synteny conservation patterns may be easier to detect with a second outgroup (notably during the threading procedure), resulting in fewer conflicts in orthology graphs and more accurate orthology constraints. To integrate information from different reference outgroups, SCORPiOs is run using each outgroup independently until the end of the graph community detection step (Supplementary Figure 1). Then, for each gene family, SCORPiOs selects the orthology/paralogy predictions from the most informative outgroup to go through the rest of the workflow. The most informative outgroup is the one that result in a larger number of gene graphs (suggesting that the other outgroup has lost one or more gene copies), or which requires the lowest number of cut edges to separate the orthogroups in the gene graph.

**Multiple WGD correction.** SCORPiOs supports the correction for several WGDs in a gene tree forest, provided that an outgroup non-duplicated species is available for each single WGD. These WGD can be independent (in different lineages) or nested (in a subclade from a more ancient WGD). When correcting multiple WGDs, SCORPiOs is run independently on each WGD to build corrected subtrees. However, duplicated species sets are handled as mutually exclusive: for instance, in trees containing both 3R and 3R+4R fish species, 4R species are excluded from 3R synteny analyses and correction. Their 3R orthology group is selected based on their closest 3R ortholog in the original tree. Corrected subtrees are regrafted in a similar manner as described in Supplementary Note 3, starting from the most recent duplications to the most ancient (tips to root). In this manner, when regrafting 3R corrected subtrees, 4R subtrees are assumed to be correct and are simply kept with the same topology and at the same position relative to their closest 3R ortholog.

**Excluded species.** SCORPiOs allows to specify duplicated species that should not be used during the pairwise synteny analyses, for instance if its genome is poorly assembled. For these species, genes will be included in the orthology table but not in the synteny orthology graphs. When building the constrained tree topologies, genes from excluded species will be placed in the same orthology group as their closest ortholog in the original gene tree.

**Parameter specifications.** Several parameters can be user-defined, such as the size of the sliding window (default=15), or adding a cut-off on synteny similarity scores to predict gene orthologies with synteny ($\Delta$S, default=0). By default, SCORPiOs also ignores tree/synteny inconsistencies when an orthogroup community contains only a single gene. These are poorly-supported WGD duplication nodes, that, in our datasets with a reasonable number of species (~10), are mostly synteny prediction errors caused by assembly or annotation artefacts. Additionally, SCORPiOs provides options to customize the relaxed orthology table: an option to optimize the minimal number of syntenic ortholog neighbours required to include an homolog in the table, and an option to ignore orthologs based on the original tree that do not have any syntenic ortholog neighbours.

| SCORPiOs ITERATION 1 Ensembl89 | | | | | | | | | | | |
|---|---|---|---|---|---|---|---|---|---|---|---|
| Families [a] | Total | Zebra-fish | Cave fish | Tetrao-don | Fugu | Stickle-back | Cod | Tilapia | Medaka | Molly | Platy-fish |
| | 15,476 | 19,737 (76 %) | 19,021 (83%) | 16,827 (86%) | 16,473 (89%) | 16,898 (81%) | 16,607 (83%) | 18,079 (84%) | 15,829 (80%) | 19,459 (82%) | 17,633 (87%) |

| Synteny orthology graphs [b] | Non-processed families | Total graphs | 2 cliques | Girvan-Newman | Kerningan-Lin |
|---|---|---|---|---|---|
| | 900 | 14,576 | 10,172 | 4,388 | 16 |

| Synteny incon-sistent trees [c] | Total | Corrected | | | Non corrected | |
|---|---|---|---|---|---|---|
| | 3,394 | Total 2,361 | ProfileNJ 1,902 | TreeBeST 459 | failed lk-test 818 | un-tested 215 |

| Likelihood-tests details [d] | Total | | ProfileNJ | | TreeBeST | |
|---|---|---|---|---|---|---|
| | equivalent | better | equivalent | better | equivalent | better |
| | 1,696 | 665 | 1,392 | 510 | 304 | 155 |

**Supplementary Table 1: Summary of SCORPiOs correction of fish subtrees in Ensembl v89 (iteration 1)**

[a] Number of teleost gene families, with the number and fraction of genes they represent in each teleost.

[b] Number of synteny orthology graphs. For 900 families, SCORPiOs did not build a graph: 771 are families with fewer than 3 genes, 66 families could not be threaded in the synteny analysis, 39 families were highly multigenic (>4 genes per species) and would not be considered for correction and 24 were in too small windows on the gar genome. We indicate the algorithm used to separate the genes in two communities (orthology groups). “2 cliques” refers to graphs where the genes were already naturally separated in two communities.

[C] Number of synteny inconsistent gene trees and subsequent corrections. For each inconsistent tree, a solution is searched with ProfileNJ first and then TreeBeST phyml if ProfileNJ fails to find a statistically equivalent tree. Non-corrected trees correspond to trees where the corrected solution was significantly worse than the original tree (failed likelihood test), or where no corrected solution was found.

[d] Likelihood test details and number of trees for which the correction yields an equivalent or statistically better supported tree (AU tests, at $\alpha = 0.05$).

| SCORPiOs ITERATION 2 Ensembl89 | | | | | | | | | | | |
|---|---|---|---|---|---|---|---|---|---|---|---|
| Families [a] | Total | Zebra-fish | Cave fish | Tetrao-don | Fugu | Stickle-back | Cod | Tilapia | Medaka | Molly | Platy-fish |
| | 15,476 | 19,731 (76 %) | 19,024 (83%) | 16,833 (86%) | 16,481 (89%) | 16,898 (81%) | 16,610 (83%) | 18,082 (84%) | 15,829 (80%) | 19,465 (82%) | 17,641 (87%) |

| Synteny orthology graphs [b] | Non-processed families | Total graphs | 2 cliques | Girvan-Newman | Kerningan-Lin |
|---|---|---|---|---|---|
| | 1,099 | 14,377 | 10,286 | 4,041 | 15 |

| Synteny incon-sistent trees [c] | Total | Corrected | | | Non corrected | |
|---|---|---|---|---|---|---|
| | | Total | ProfileNJ | TreeBeST | failed lk-test | un-tested |
| | 1,028 | 35 | 18 | 17 | 784 | 209 |

| Likelihood-tests details [d] | Total | | ProfileNJ | | TreeBeST | |
|---|---|---|---|---|---|---|
| | equivalent | better | equivalent | better | equivalent | better |
| | 1,696 | 665 | 1,392 | 510 | 304 | 155 |

**Supplementary Table 2: Summary of SCORPiOs correction of fish subtrees in Ensembl v89 (iteration 2)**

[a] Number of teleost families, as in Table S1.

[b] Number of synteny orthology graphs, as in Table S1. For 1099 families, SCORPiOs did not build a graph: 771 are families with less than 3 genes, 206 fell in window with non-updated synteny context compared to iteration 1, 63 families could not be threaded in the synteny analysis, 35 families were highly multigenic (>4 genes per species) and 24 were in too small windows on the gar genome.

[C] Number of synteny inconsistent gene trees, as in Table S1.

[d] Likelihood test details, as in Table S1.

Supplementary Table 3 : Gene ontology (Biological Process) enrichment results for zebrafish genes retained as duplicates in all species after the TGD. [separate .xls file]

Supplementary Table 4 : KEGG pathway enrichment results for zebrafish genes retained as duplicates in all species after the TGD. [separate .xls file]

Supplementary Table 5 : Gene ontology (Biological Process) enrichment results for zebrafish genes retained as duplicates in some species after the TGD. [separate .xls file]

Supplementary Table 6 : KEGG pathway enrichment results for zebrafish genes retained as duplicates in some species after the TGD. [separate .xls file]

Supplementary Table 7 : Gene ontology (Biological Process) enrichment results for zebrafish genes returned to singletons in all species after the TGD. [separate .xls file]

Supplementary Table 8 : KEGG pathway enrichment results for zebrafish genes returned to singletons in all species after the TGD. [separate .xls file]

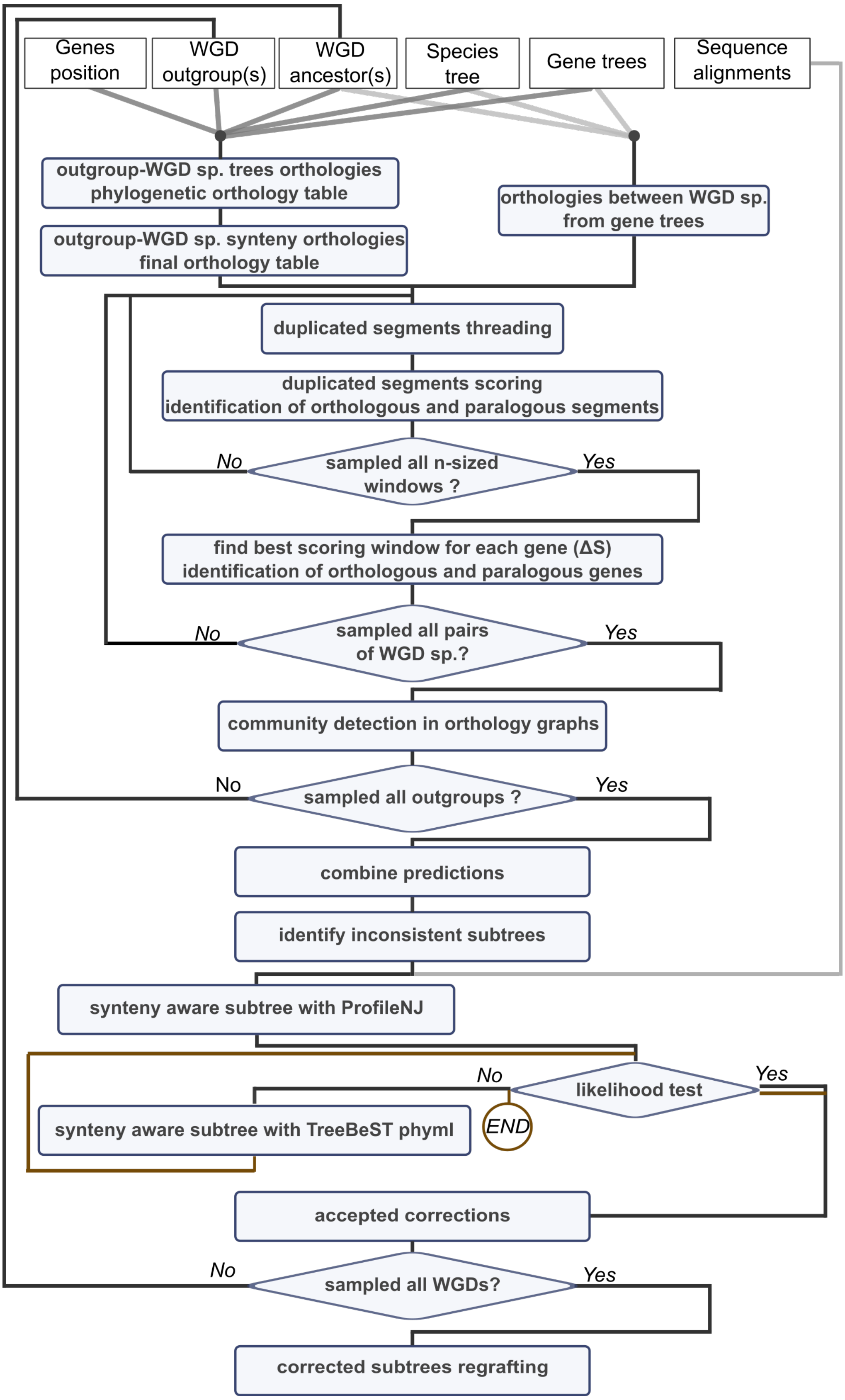


**Supplementary Figure S1: SCORPiOs workflow.** Flowchart of the SCORPiOs workflow to correct an input gene tree forest containing species that underwent one or several WGDs, using one or several unduplicated outgroups. See main text and Supplementary notes for details.

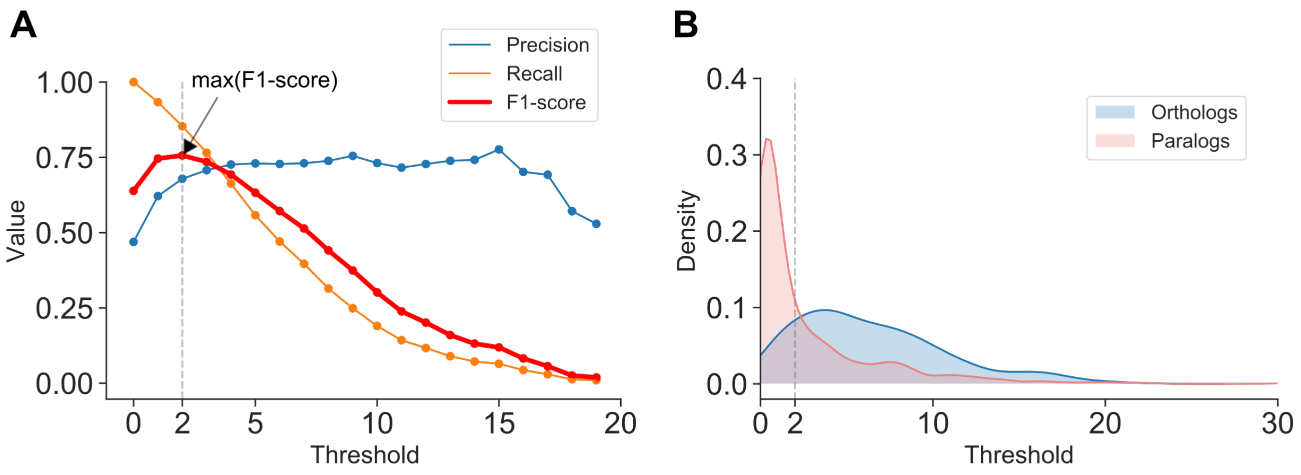


**Supplementary Figure S2: Recovery of putative orthologs based on their syntenic context. A.** We collected teleost orthologs and paralogs of spotted gar genes based on the original gene forest, and determined the optimal number of ortholog syntenic neighbours to differentiate orthologs from paralogs. We used the F1-score to combine Precision and Recall, which was maximal at a threshold of 2 syntenic orthologs for this dataset. **B.** Distributions of the average number of syntenic orthologs. The threshold (dashed line) excludes the largest fraction of paralogs. At this threshold, only 32% of predicted orthologs are false positives, and we recover over 85% of all true orthologs.

This threshold can be optimized in SCORPiOs by setting the optional argument `optimize_synteny_support_threshold` to `y` in the configuration file, as synteny conservation and genome assembly contiguity may impact the optimal threshold for other datasets.

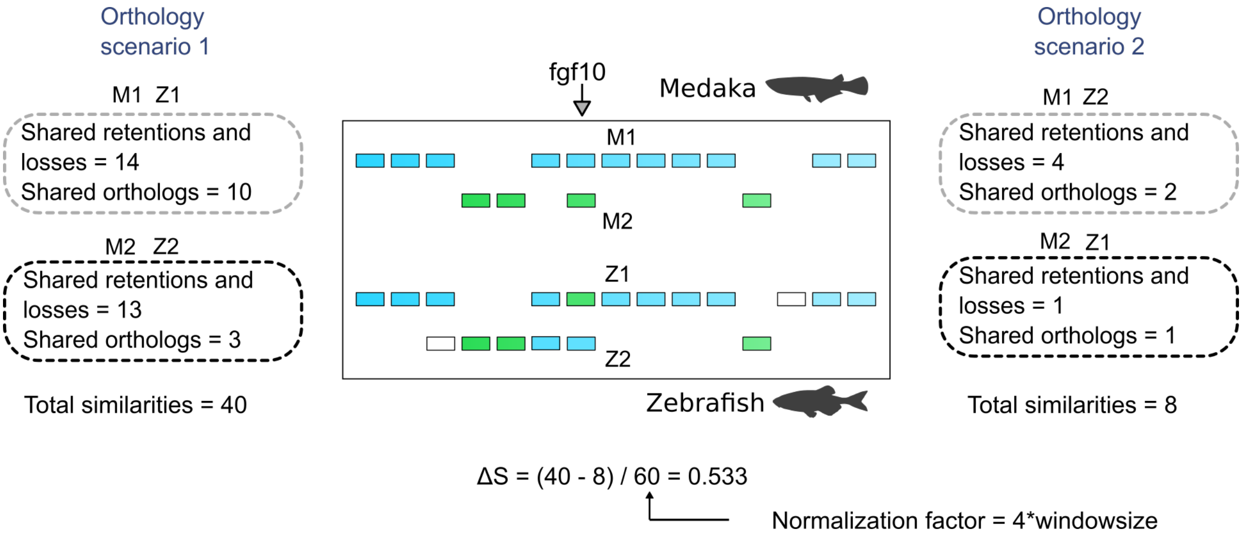


**Supplementary Figure S3: Example for the computation of a ΔS score between threaded segments of two duplicated species.** First, the similarity of threaded segments in each of the two orthology scenarios is assessed using the number of shared gene retentions/losses and annotated orthologs. Second, the resulting similarity scores are compared between scenarios and normalized to obtain the ΔS score.

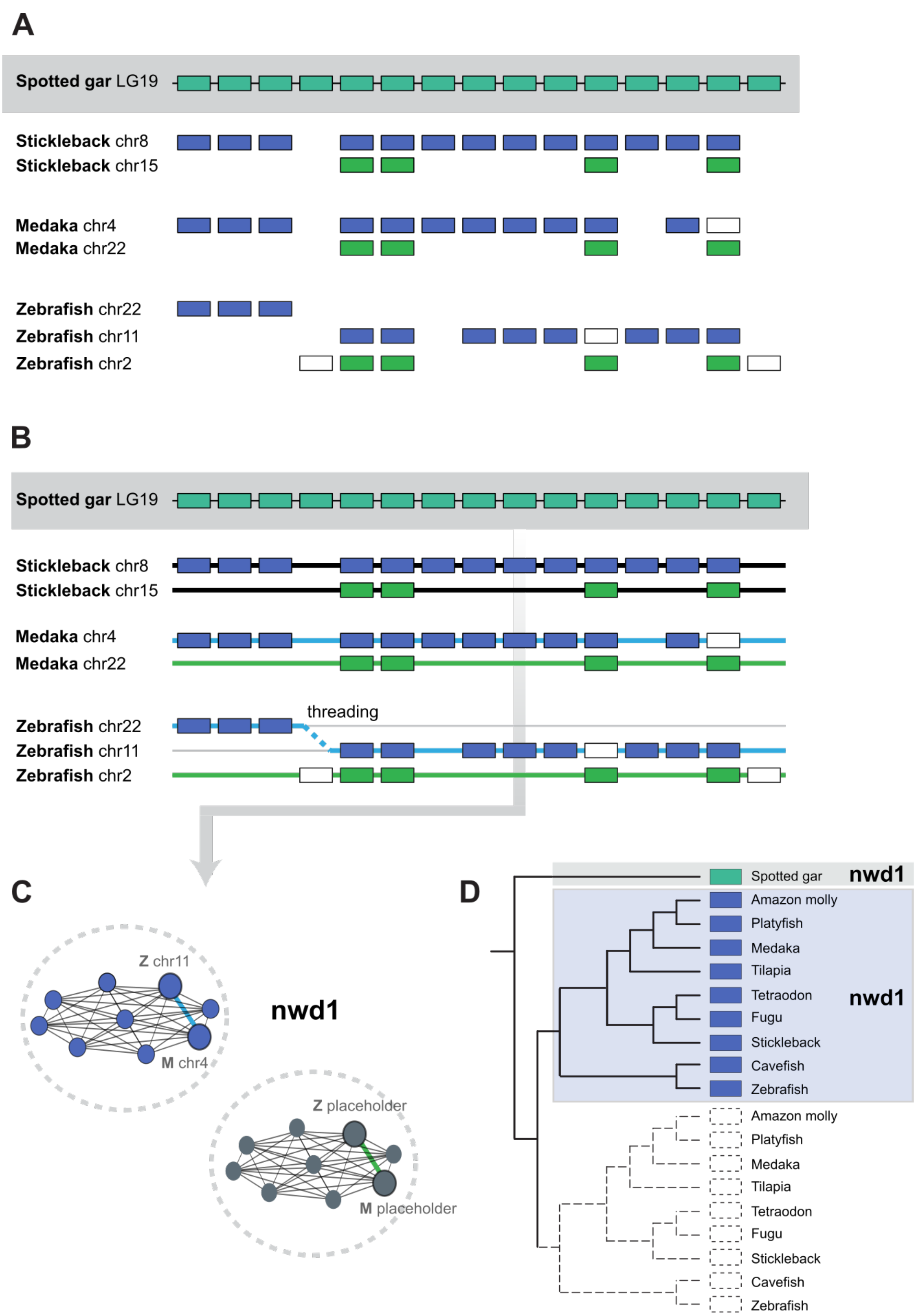


**Supplementary Figure S4**. Example of a SCORPiOs tree for a gene family (*nwd1*) where only one WGD gene copy was retained in all teleost species (early gene loss). The orthology graph still contains two communities, as gene losses are explicitly accounted for by placeholder nodes (as shown in C).

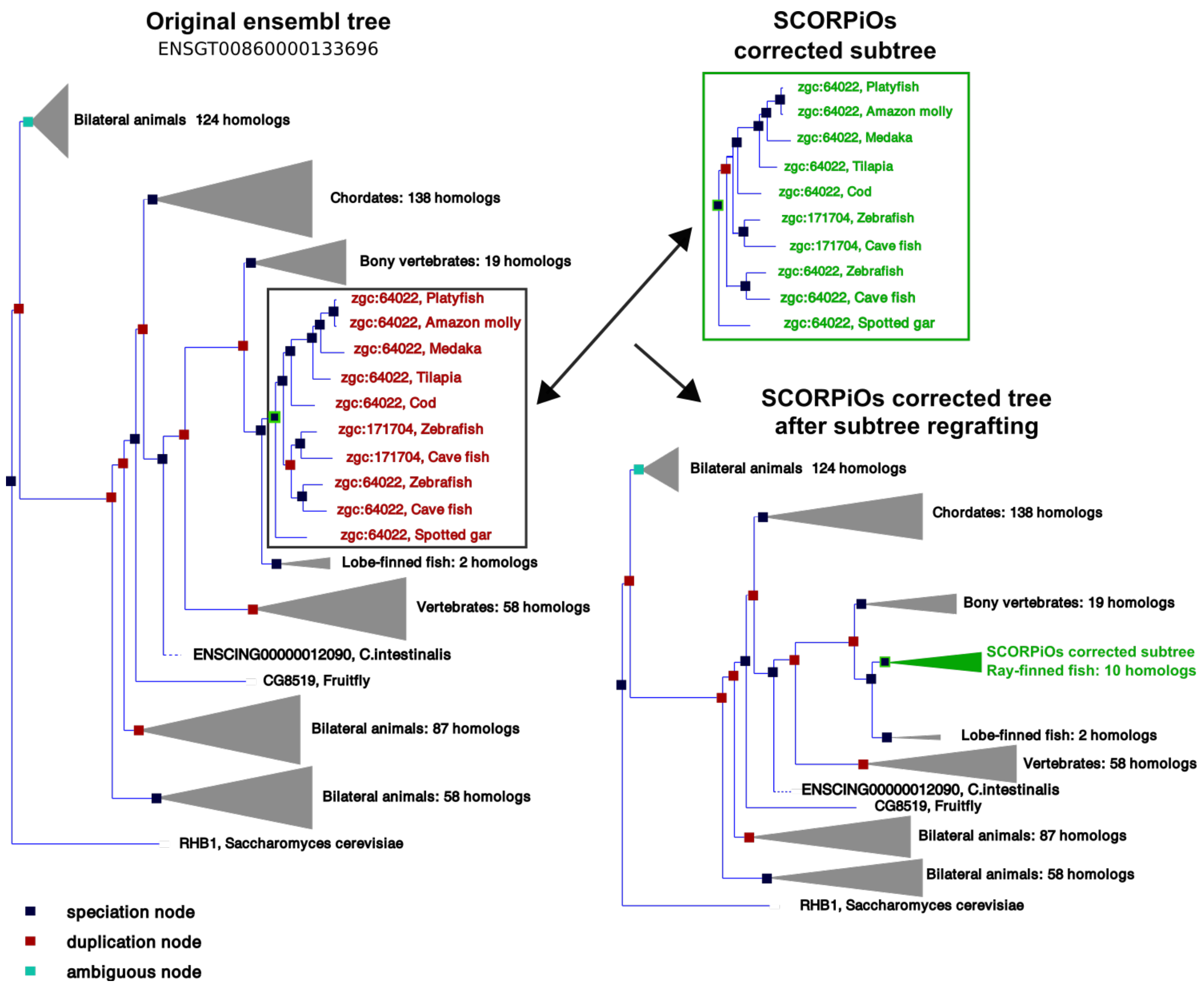


**Supplementary Figure S5: Example of SCORPiOs-corrected subtree regrafting into the original gene tree.** Genes from the corrected subtree form a clade in the original gene tree. Regrafting the subtree simply consist in removing the original subtree and grafting the corrected version in its place. SCORPiOs then recomputes branch lengths for the full tree.

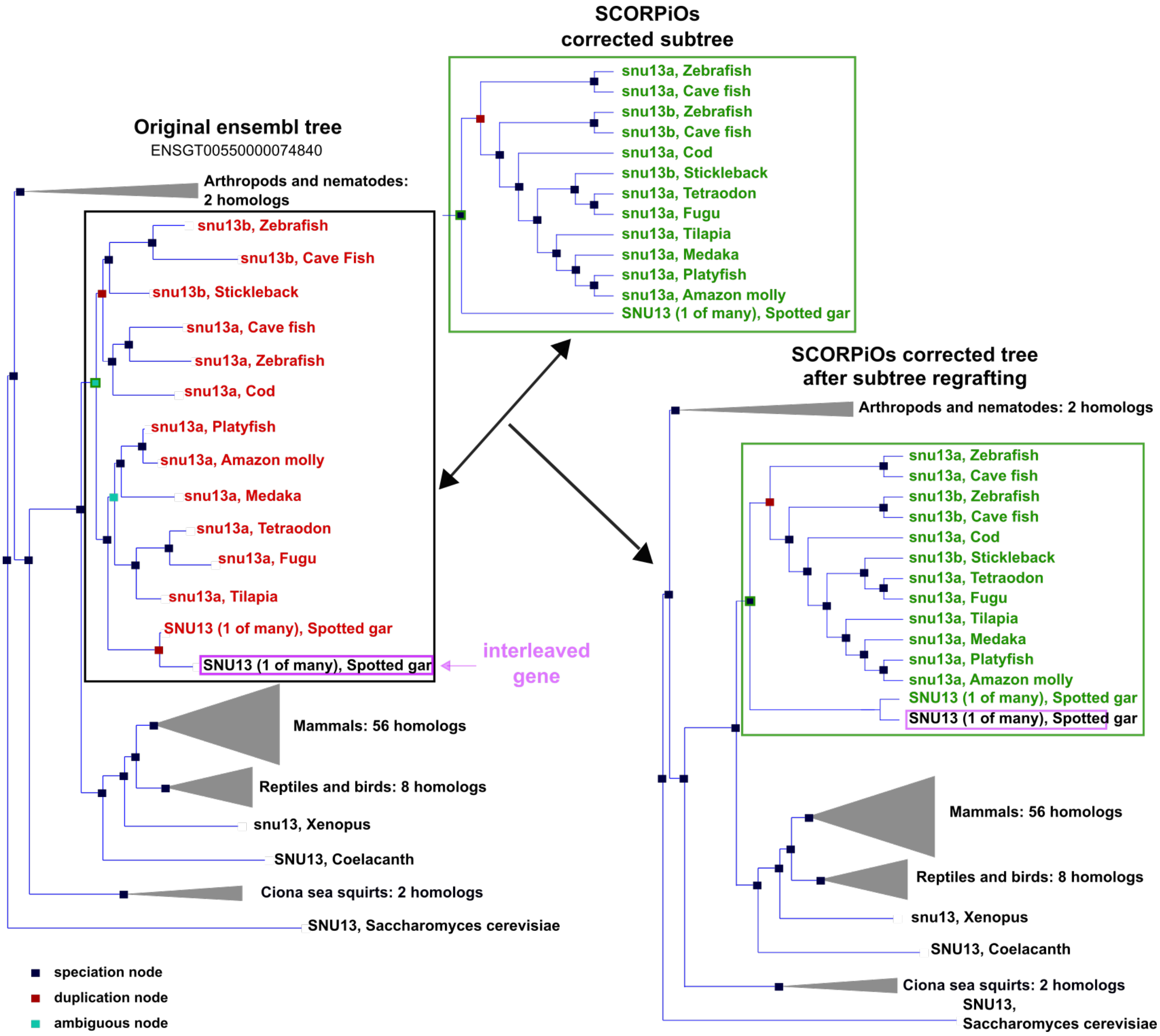


**Supplementary Figure S6: Example of SCORPiOs corrected subtree regrafting into the original gene tree.** Genes of the corrected subtree do not form a complete clade in the original gene tree. A second gene copy of the spotted gar is present (interleaved gene). However, the position of this gene is consistent with the species tree, so SCORPiOs leaves its position unchanged in the resulting corrected full tree topology. SCORPiOs then recomputes branch lengths for the full corrected tree.

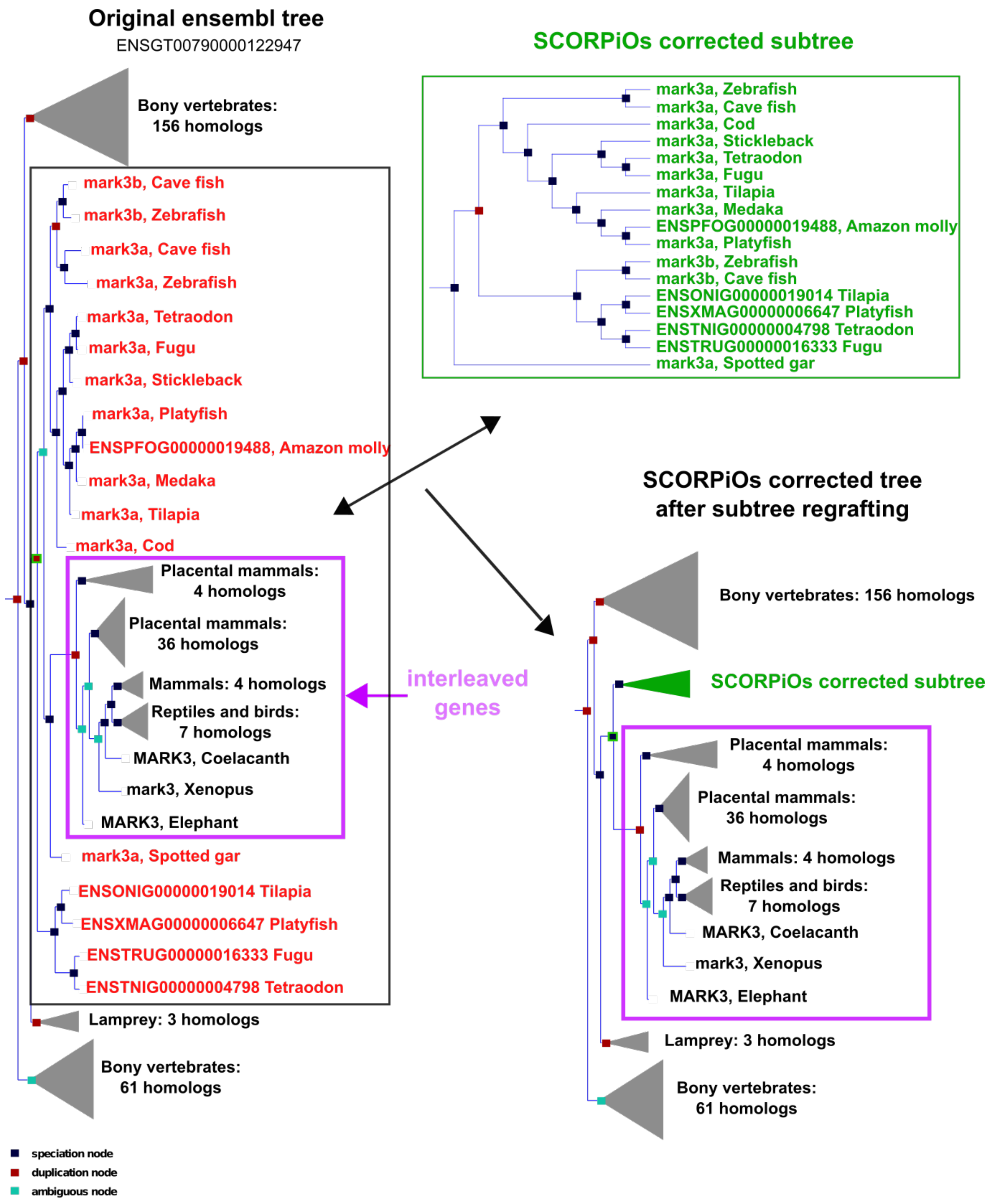


**Supplementary Figure S7: Example of SCORPiOs corrected subtree regrafting into the original gene tree.** Genes of the corrected subtree are not together in a clade in the original gene tree. Mammals and birds genes are interleaved in the original sub-tree. As these genes do not descend from the ancestral unduplicated Ray-finned fish gene, SCORPiOs places them as outgroup to the corrected sub-tree, leaving their topology unchanged in the resulting corrected full tree topology. SCORPiOs then recomputes branch lengths for the full corrected tree.

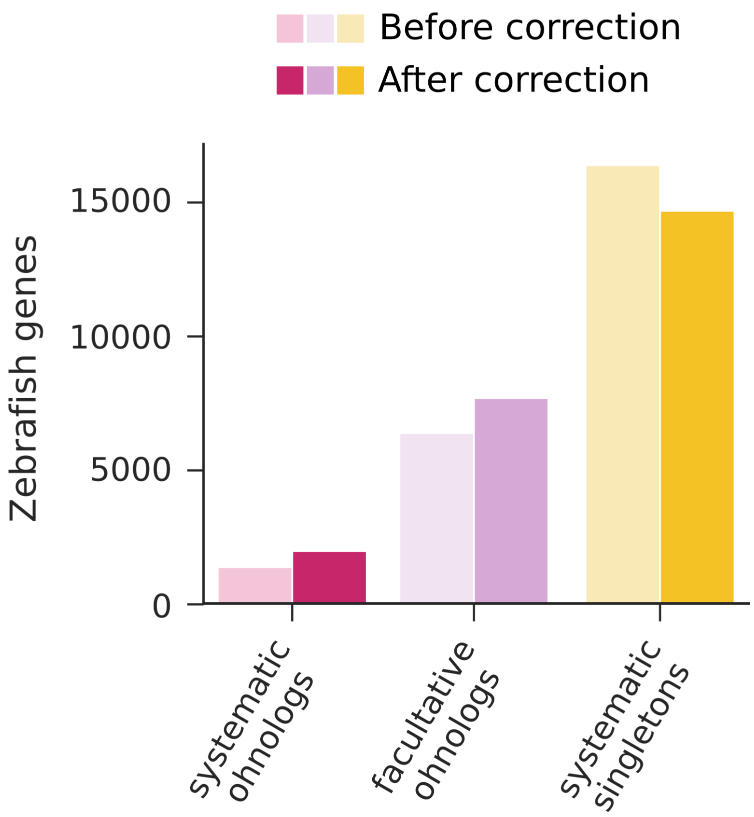


**Supplementary Figure S8: Evolutionary categorization of zebrafish genes.** Counts of systematic ohnologs, facultative ohnologs and systematic singleton using the Ensembl gene tree forest (before correction) and the SCORPiOs-corrected forest (after correction).

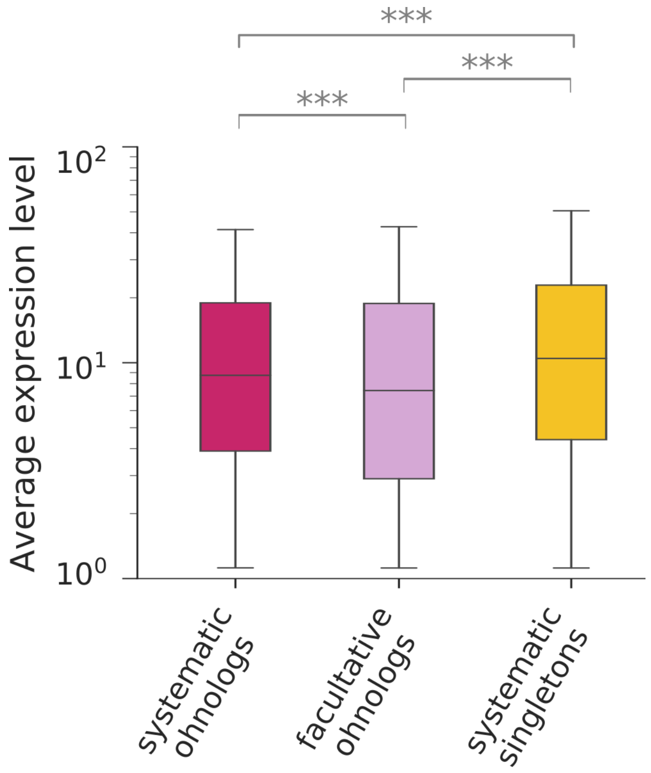


**Supplementary Figure S9**: **Average expression (quantile normalised TPM) in zebrafish of genes with different evolutionary trajectories after the TGD.** Expression levels are compared across categories using Wilcoxon–Mann–Whitney tests (*** $p < 10^{-3}$).
